# Phylobioactive hotspots identified through multidimensional profiling of botanical drugs used to treat Chagas disease in Bolivia and Dioscorides’ De Materia Medica

**DOI:** 10.1101/862029

**Authors:** Andrea Salm, Sandhya R. Krishnan, Marta Collu, Ombeline Danton, Matthias Hamburger, Marco Leonti, Giovanna Almanza, Jürg Gertsch

**Author notes:** authors contributed equally to the study (Andrea Salm, Sandhya R. Krishnan).

## Abstract

Globally, more than six million people are infected with *Trypanosoma cruzi*, the causative protozoan parasite of the vector-borne Chagas disease (CD). In Bolivia, CD is hyperendemic and a major health problem among indigenous communities. Although botanical drugs are used widely among different ethnic groups in Bolivia, studies challenging the hypothesis that effective antitrypanosomal medicinal agents were identified empirically are lacking. We conducted a cross-sectional ethnopharmacological field study in Bolivia among different ethnic groups in the Chaco, Chiquitanía and Inter-Andean valleys. We compared botanical drugs used in Bolivia in the context of CD with botanical drugs from unrelated indications from the Mediterranean *De Materia Medic*a (*DMM*) compiled by Dioscorides two thousand years ago. A total of 775 ethyl acetate plant extracts with and without ethnomedical indications for CD treatment were profiled against *T. cruzi* epimastigote and procyclic *T. brucei* proliferation, parasite release from *T. cruzi* trypomastigote infected cells, as well as for host cell cytotoxicity *in vitro*. Inhibition of parasite release was monitored using a flow cytometry-based celluar assay. At 25 µg/mL, less than 5% of all extracts exhibited selective toxicity for *T. cruzi*. We found no evidence that ethnomedicine-inspired bioprospecting significantly increased the probability of finding selective antichagasic botanical drugs. The ethnomedical data further indicate a discrepancy between local and scientific concepts about CD among the studied ethnic groups. Intriguingly, the phylobioactive anthraquinone hotspot identified in this study matched the antichagasic activity of *Senna chloroclada*, the taxon with the strongest consensus for treating CD among the Izoceño-Guaraní. Selected antitrypanosomal plant extracts from *DMM* were subjected to HPLC-based activity profiling and targeted isolation of active compounds yielding sesquiterpene lactones, naphtoquinones and anthraquinones. Because the anthraquinone emodin selectively and potently inhibited *T. cruzi* in host cell infection, we performed a preliminary structure-activity relationship analysis for the 9,10-anthracenedione scaffold, exploring the impact of differential hydroxylation. This study shows that the multidimensional phylobioactivity-guided identification of antichagasic natural products enables comparative bioprospecting and is suitable to challenge ethnopharmacological hypotheses.

**Author summary:** Chagas disease (CD) is a parasitic disease caused by the protozoan *Trypanosoma cruzi*. In Bolivia, CD is a major health problem among indigenous communities, which frequently use traditional medicine to treat the chronic symptoms of the disease related to cardiomyopathy. However, the ethnomedical context of the use of such remedies is largely unclear and it remains unknown whether the botanical drugs have any effect on parasitemia. In a field study among different ethnic groups in the Chaco, Chiquitanía and Inter-Andean valleys the authors collected ethnobotanical and ethnopharmacological information. Later, they profiled and compared the CD botanical drug extract library from Bolivia with a botanical drug extract library from the Mediterranean *De Materia Medic*a with no association to CD. Using phylogenetic and biological information, they identified bioactive hotspots among different taxa and isolated antichagasic natural products. This led to a first structure-activity relationship study of the natural product class called anthraquinones. While there was no overall statistical difference between the libraries, it is noteworthy that the botanical drug derived from *Senna chloroclada* with the highest consensus among the Guaraní communities, also belonged to the anthraquinone cluster, potentially providing a molecular explanation for its use.

## Introduction

Chagas disease (CD) or American trypanosomiasis is the disease caused by infection of the haemoflagellate protozoan *Trypanosoma cruzi* (1). The parasitic *T. cruzi* is mainly transmitted to humans by hematophagous Reduviid bugs of the subfamily Triatominae (*Triatoma* spp.) (1–3). Other means of transmission involve blood transfusion, organ transplant, congenitally, and oral contamination (1, 2). CD is classified as one of the most neglected tropical diseases, especially among low-income populations (4). The WHO estimates that about 6 to 7 million people are infected worldwide, primarily in Latin America, where the disease is endemic (5). Bolivia has the highest CD incidence in the world (6, 7), and *T. cruzi* infection has a high prevalence in different rural areas, primarily affecting the indigenous communities in the Inter-Andean valleys and the Chaco (7–9).

CD presents two clinical phases. The acute stage of the disease (i.e. infection) remains often unperceived and only very rarely morbidity and mortality occur (10). In some cases, the sites of bug bites get inflamed and form nodules called chagoma or “Romaña sign” when the protozoan enters via the conjunctiva. After an incubation period of less than two weeks, newly infected individuals may develop fever, chills, myalgia, rash or meningeal irritation. However, the initial chagoma can remain the only indication of *T. cruzi* infection for years. At the chronic stage, parasitism and inflammation of the heart and/or enlarged colon can result in severe pathophysiological endpoints, including megacolon, megaoesophagus and cardiomyopathy (11). Cardiomyopathy manifests as arrhythmias, congestive heart failure or sudden cardiac arrest. During the chronic phase, *T. cruzi* infection can present an indeterminate form, where, depending on the *T. cruzi* strain and possibly host immunity, approximately 70% of the infected individuals remain completely asymptomatic (2,3,12). Currently, there are two approved drugs that are used to treat acute CD, namely the nitroheterocyclic compounds nifurtimox (Lampit^®^) and benznidazole (Rochagan^®^). However, there is a controversy about their efficacy in chronic CD (13, 14) and both drugs show adverse effects upon prolonged administration (15, 16). Moreover, these antichagasic drugs are hardly accessible for the indigenous communities in Bolivia. Consequently, people living in rural areas of Bolivia rely on traditional medicines for their primary health care (17, 18). Botanical drugs contribute significantly to Bolivian folk medicine and they are harvested wild, cultivated and traded at local markets (19). Despite reports making general claims about the efficacy of traditional botanical drugs (20), few studies address their mode of action based on comparative hypothesis-driven ethnopharmacological research. It is unknown whether CD, which has a millennial history in the area (21), enforced a selection pressure to prompt ethnomedical strategies among the affected indigenous groups in Bolivia that directly reduce parasitemia or primarily treat the symptoms of CD.

Plants are known to produce a high diversity of secondary metabolites showing numerous pharmacological effects, including antiparasitic and antimicrobial (22, 23). Medicinal plants have been shown to act via different mechanisms of action also against various causative agents of neglected tropical diseases like *Mycobacterium ulcerans* or *T. cruzi* (24–26). Plant secondary metabolites have led to the development of antiprotozoal chemotherapeutics, which include antimalarials such as the alkaloid quinine from *Cinchona* spp. (Rubiaceae) and the sesquiterpene lactone (SL) artemisinin from *Artemisia annua* L. (Asteraceae) (27). Ethnopharmacology inspires bioprospecting as it investigates treatment consensus of natural drugs among ethnic groups as anecdotal indications for presumed pharmacological efficacy (28, 29). Considering Bolivia’s rich biocultural diversity and the endemic nature of CD, ethnopharmacological and bioprospecting studies focusing on antichagasic remedies are surprisingly scarce (30–33). In parallel, few studies were dedicated to the ethnomedicine and management of CD in Bolivia (34, 35). Although different plant extracts and natural products have been reported to exert moderate (IC_50_ values ≤ 100 µg/mL) to significant (15 ≤ µg/mL) selective toxicity towards *T. cruzi* epimastigotes (36–39), reports on activity against trypomastigote cellular infection and amastigote replication *in vitro* are relatively rare. Overall, systematic and comparative antitrypanosomal screenings with comprehensive extract libraries derived from botanical drugs are lacking, thus hindering a direct comparison of efficacy.

The aim of the present study was to conduct a cross-sectional ethnopharmacological study among the indigenous groups Quechua, Izoceño-Guaraní, Ayoreo and Chiquitano to investigate their prevalent ethnomedical strategies to treat CD in Bolivia. This included the documentation of the knowledge about botanical drugs used in treating CD-related symptoms. Fieldwork involved structured and semi-structured interviews, plant collections (drug samples and herbarium vouchers), and identification of the plant taxa used. An ethyl acetate (EtOAc) extract library of the botanical drugs used to treat CD was generated for *in vitro* testing against trypanosomes. To validate the ethno-directed bioprospecting approach, we also profiled *in vitro* antitrypanosomal effects of EtOAc extracts obtained from Mediterranean botanical drugs described in *De Materia Medica* (*DMM*), written by Dioscorides in the first century AD (40). We hypothesized that the CD botanical drug library would result in a comparatively higher number of extracts showing selective antichagasic activity *in vitro*. Using phylobioactivity-guided profiling in combination with HPLC-based activity profiling of active extracts we tested whether bioactive phylogenetic clusters can serve as a basis for the more efficient characterization of antichagasic plant metabolites. The *DMM* extract library yielded phylogenetic patterns of activity and different active hotspots could be identified representing chemotaxonomic associations. This multidimensional profiling resulted in a first preliminary structure-activity relationship (SAR) study on the antitrypanosomal effects of anthraquinones on parasite release upon host cell infection by trypomastigotes *in vitro*.

## Methods

### Ethnopharmacological fieldwork

The study was carried out among four indigenous groups (Quechua, Izoceño-Guaraní, Chiquitano and Ayoreo), settled in geographically distinct rural areas in Bolivia (Fig. 1). The surveyed Quechua communities were located in the Inter-Andean valleys between 3000 and 2200 m.a.s.l. in the municipality of Mizque (Cochabamba Department). The Guaraní in Bolivia live in the Chaco region, which is a semi-arid plain grassland, interspersed with swamps and thorny forests, that extends across southeast lowland Bolivia. In Bolivia, there are three subgroups of Guaraní (Ava, Simba and Izoceño), marked by linguistic and historical differences. We visited communities of the Izoceño-Guaraní in the municipality of Charagua (Santa Cruz Department). The Chiquitano informants lived in the lowland Chiquitanía region situated in the Santa Cruz Department. This region is an ecologically transitional zone between the arid plains of the Chaco and the tropical rain forests. Today, Chiquitanos are native Spanish speakers. The survey was carried out in the Chiquitano communities in the municipalities of Concepción, San José de Chiquitos and Roboré. The Ayoreo groups used to live in the Chaco but have been sedentarized and acculturated by missionaries in the mid twentieth century. We surveyed Ayoreo communities situated in the Chiquitanía region (municipalities of Concepción, Pailón and San José de Chiquitos) and Ayoreo informants who lived in the city of Santa Cruz. All visited indigenous communities were small settlements located in rural areas characterized by a low population density. The main economic activity was subsistence farming. Traditional dwellings were predominantly constructed of adobe walls, earthen floors and thatched roofs, and constitute an ideal habitat for the CD vector *T. infestans* (41). *T. cruzi* transmission was hyperendemic showing the highest infection rates in the Chaco region and Inter-Andean valleys (42–46). Biomedical health care was mainly provided by primary health centers located in the bigger villages Charagua, La Brecha and Iyovi (Chaco), San José de Chiquitos, Roboré and Concepción (Chiquitanía), Mizque and Laguna Grande (Inter-Andean valleys). However, most of them were insufficiently equipped, and CD treatment was marginally successful (41). Additionally, a survey was conducted in the herbal markets of the cities of La Paz and Santa Cruz de la Sierra.

**Figure 1.**
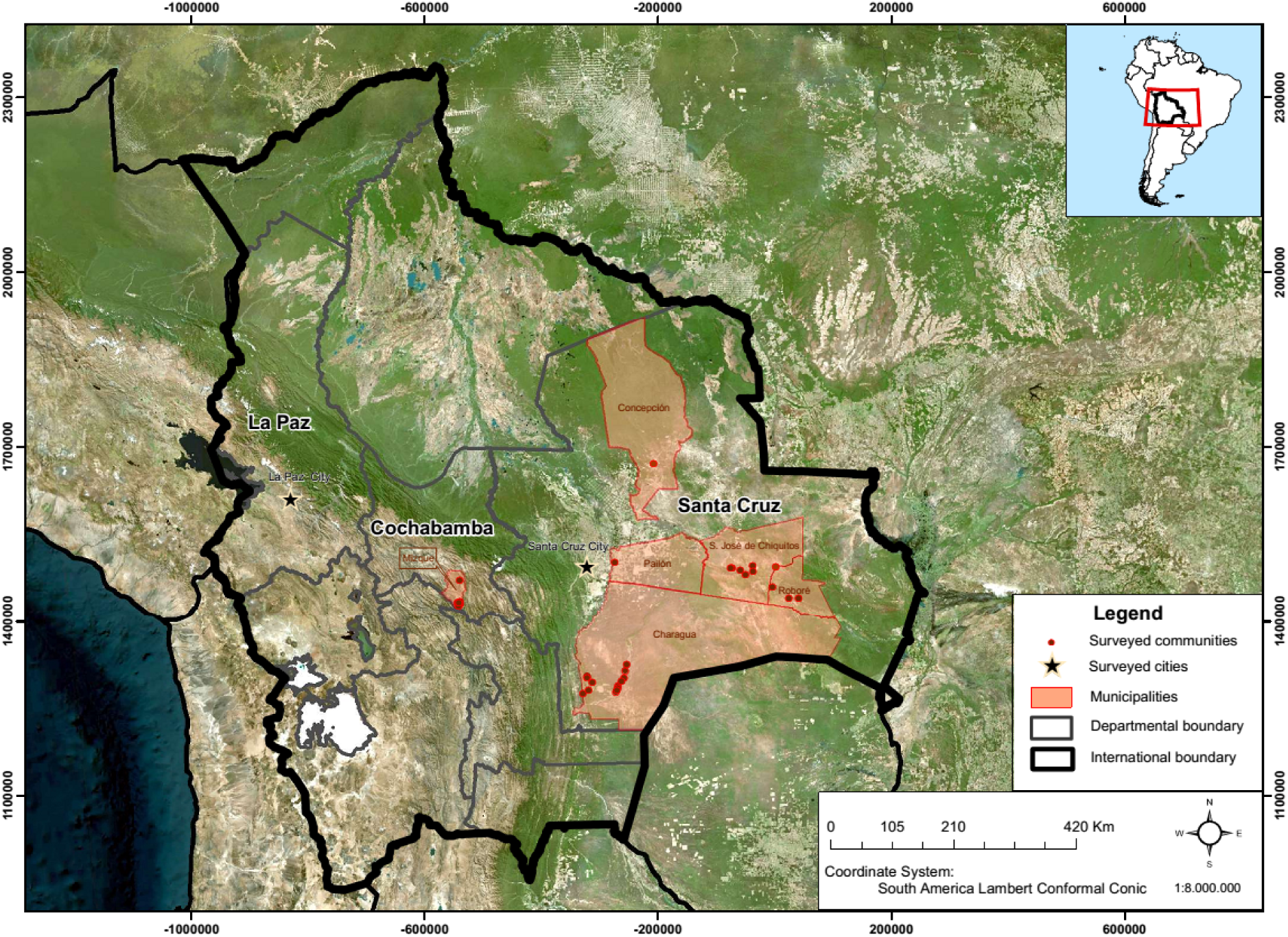
Areas of the ethnopharmacological field study in Bolivia. The three different municipalities (orange), surveyed communities (red dots) and cities (stars) are shown. The map was generated using the GPS coordinates with ArcGIS, version 10.3.

Fieldwork was carried out between June 2014 and May 2015 in all study sites during all vegetation periods. The survey was directed towards the management of CD related symptoms and associated knowledge. Before starting with the survey, the project was presented to the local authorities and to the communities (Fig. 1) during initial meetings for approval. Local research participants were randomly selected according to their availability, willingness and individual interest while each participant represented a distinct household. Interviews took place at the participants’ home and were conducted in Spanish, or in their native language (Guaraní, Ayoreo, Quechua) with the assistance of a local translator. In the cities, we interviewed the medicinal plant vendors at the herbal markets from where we directly purchased the botanical drugs. Ethnomedical and ethnopharmacological information was obtained through structured and semi-structured interviews, free listing of remedies, observing practices *in situ* and by collecting plant material during walk-in-the-woods together with the participants. During interviews, representative photographs of chagoma were shown and symptoms of chronic CD (cardiomyopathy, megacolon, megaoesophagus) described to the participants for eliciting focused responses. Questions targeted ethnomedical concepts related to CD and plant taxa used for treatment of acute and chronic symptoms. Vernacular plant names, parts used, dosage, mode of preparation and administration, provenance and availability were documented. Botanical voucher specimens were collected with the help of local informants, prepared and deposited at the National Herbarium in Bolivia (LPB) under the collection ID ASMPx. Botanical identification was carried out by specialists at LPB, and botanical names standardized in accordance with the theplantlist.org (47) and plant families following Angiosperm Phylogeny Group 4 (APG IV, 2016) (48). Data were analyzed with basic descriptive statistics such as the quantification of individual use reports cited by the informants according to current standards and recommendations (49).

### Ethics statement

Field research was approved by the Ministerio de Ambiente y Desarrollo Sostenible through a collaboration with the Instituto de Investigaciones Quimicas The institutional review board (IRB) of the Universidad Mayor de San Andres (UMSA FCPN) as part of the National Bioethics Committee approved the study. Oral consent was obtained from each research participant for treating personal data and the recording of ethnomedical information. Community consent was documented on video or tape. Written consent at the time of the study was not feasible as the majority of participants were illiterate.

### Collection of botanical drugs mentioned in De Materia Medica

Botanical drugs mentioned in *DMM* and associated voucher specimens were collected in several locations in Europe and the Mediterranean area, cultivated in domestic gardens or purchased from commercial suppliers between 2014 and 2016. Plant taxa were identified using the Flora Europea (50). Only those 660 botanical drugs, whose botanical description in *DMM* permitted a taxonomic identification (40) were collected. Prior to solvent extraction samples were dried at 40–60 °C. Herbarium voucher specimens were identified at the Department of Biomedical Science (University of Cagliari) and voucher specimens deposited at the Herbarium of the Botanical Garden of Geneva (Switzerland) and at the Herbarium of the National and Kapodistrian University of Athens (ATHU).

### Preparation of extracts

Medicinal plant parts used in the context of CD were collected *in situ* and from the same collections herbarium voucher specimens were prepared. The clean plant materials (roots washed) were air-dried in the shade and stored dry in cotton bags at −20 °C prior to extraction. The extracts were generated as follows: Air-dried and powdered (0.5 mm) plant material was used for the preparation of semipolar extracts. The samples were exhaustively extracted by maceration with EtOAc (99.8%, Sigma Aldrich) at room temperature for 48 h. EtOAc (polarity index P = 4.4) was used based on its optimal extraction for secondary metabolites that are apolar to semipolar, excluding the highly polar constituents like sugars and polymers. Upon filtration (Whatman, 2.5 µm), samples were concentrated under reduced pressure on a rotary evaporator. The dried extracts were used to prepare stock solutions of 5 mg/mL in dry DMSO (≥99%, Sigma Aldrich) for subsequent use in *in vitro* assays. Crude extracts and DMSO stock solutions were stored at −20 °C.

### Biological assays

#### In vitro activity against T. cruzi epimastigotes

Antitrypanosomal activity was first evaluated against epimastigotes of *T. cruzi* (Y strain obtained from ATCC) by the XTT assay (51, 52). Since all known antichagasic agents show toxicity also against the epimastigote form, we used this screening as a preselective assay. Parasites were maintained in liver infusion tryptose medium (LIT) supplemented with 10% FBS (fetal bovine serum) at 27 °C, prepared according to the recommendations by ATCC. Epimastigotes in exponential growth phase were counted, adjusted to a concentration of 1.5 x 10^6^ parasites/mL, and exposed to 6 concentrations of each plant extract (in DMSO) ranging from 100 μg/mL to 0.8 μg/mL for 72 h at 27 °C. For the *DMM* library we chose a concentration of 25 µg/mL based on recommendations for the pure compounds (53). The assays were carried out in 96-well plates (200 μL/well). After 72 h of incubation, the plates were inspected under an inverted microscope to assure growth of the controls and sterile conditions. 50 μL of a XTT (Sigma Aldrich, MO, USA)/PMS (phenazine methosulfate, Sigma Aldrich, MO, USA) solution (0.5 mg XTT/0.025 mg PMS/mL in PBS) were added to the plates. The parasites were further incubated at 27 °C for 3 h. Then, methanol (50 μL/well) was added, and the plates were incubated for 10 min to fix the parasites. Absorbance was determined spectrophotometrically at 490 nm on a Tecan plate reader. DMSO was used as negative control, and benznidazole and nifurtimox (both Sigma Aldrich, Switzerland) were used as positive controls. Results were expressed as IC_50_ and calculated by GraphPad Prism 5^®^. Each assay was carried out at least in three independent experiments each in triplicate.

#### In vitro activity against procyclic T. brucei brucei

The *T. b. brucei* procyclic form (427 strain) was cultured at 27 °C in SDM79 medium, supplemented with 5% FBS and hemin, as reported (54). Parasites in exponential growth phase were counted, adjusted to a concentration of 1.0 x 10^6^ parasites/mL and exposed to plant extracts for 72 h at 27 °C. The assays were carried out in 96-well plates (100 μL/well). After 72 h of incubation, the plates were inspected under an inverted microscope to enusre growth of the controls and sterile conditions. 10 μL MTT (5 mg/mL in PBS) were added to the plates. The cells were further incubated for 4 hours at 27 °C. After incubation the purple formazan precipitates were solubilized in DMSO (200 μL/well). Absorbance was determined spectrophotometrically at 550 nm. DMSO was used a negative, and nifurtimox as positive control. Results were expressed as IC_50_ calculated by GraphPad Prism 5^®^ from the sigmoidal concentration-response curve. Each assay was carried out in triplicate for the initial profiling.

#### Cytotoxicity measurements in HeLa, CHO and Raw264.7 cells

Cytotoxicity/antiproliferative assays were performed by the MTT method as reported previously (55). Immortalized hamster ovary CHO-K1 (ATCC^®^ CCL-61™), human cervical HeLa (ATCC^®^ CCL-2™) and mouse macrophage Raw264.7 (ATCC^®^ TIB-71™) cells were cultured in RPMI-1640 medium supplemented with 10% FBS at 37 °C in 5% CO_2_ atmosphere. Cells were counted and adjusted to a final concentration of 2×10^4^ cells/mL. 100 μL were seeded in 96-well plates and incubated for 24 h at 37 °C with 5% CO_2_. Then, 100 μL medium containing plant extracts at various concentrations (from 100 μg/mL to 0.8 μg/mL) were added. Plates were again incubated for 72 h at 37 °C with 5% CO_2_. After incubation, the plates were inspected under an inverted microscope to assure growth of the controls and sterile conditions. Medium was removed and 90 μL fresh culture medium and 10 μL MTT (5 mg/mL in PBS) were added to the each well. Plates were incubated for 4 hours at 37 °C with 5% CO_2_. Subsequently, the purple formazan precipitate was solubilized in DMSO (200 μL/well) and absorbance determined spectrophotometrically at 550 nm. Cells cultured in absence of compounds were used as control of viability (negative control). Results were expressed as IC_50_ or as % cell viability of negative control calculated with GraphPad Prism 5^®^. Each assay was carried out in triplicate in three independent experiments.

#### Parasite release from trypomastigote-infected cells using flow cytometry

The EtOAc extracts from the CD library and the active antitrypanosomal extracts from the *DMM* library were evaluated in an *in vitro* model of *T. cruzi* parasite relase assay in CHO cells using analytical fluorescence activated cell scanning (FACS). In this assay, it was possible to simultaneously assess inhibition of growth and release of the parasite from infected host cells, which are relevant disease parameters. The host cells were cultured in RPMI-1640 medium supplemented with 10% FBS at 37 °C in 5% CO_2_ atmosphere. Metacyclogenesis of epimastigotes (strain Y) was induced by Grace insect medium (Sigma Aldrich, MO, USA) supplemented with 10% FBS and hemin (25 µg/mL, Sigma Aldrich, MO, USA). Trypomastigotes were generated by infecting CHO cells with metacyclic trypomastigotes harvested after 9 days of cultivation in Grace medium (56, 57). Trypomastigotes of *T. cruzi* were obtained from the extracellular medium of infected CHO cells at day 5 post infection. CHO cells were counted and adjusted to a final concentration of 8×10^4^ cells/mL. 500 μL were seeded in 24-well plates and incubated for 24 h at 37 °C with 5% CO_2_. Subsequently, cells were infected with trypomastigotes in fresh medium without FBS at a multiplicity of infection (MOI) of 1:10 and incubated at 37 °C in 5% CO_2_ for 24 h. During this time, trypomastigotes were allowed to invade host cells. After infection, the medium containing non-internalized parasites was removed and cells were washed twice with PBS. 500 μL of fresh RPMI medium without FBS containing plant extracts or controls (vehicle or benznidazole) were added. Plates were then incubated for 5 days at 37 °C with 5% CO_2_. After incubation, the plates were inspected under an inverted microscope to assure cell viability and sterile conditions. To determine the antitrypanosomal effects, trypomastigotes (and residual amastigotes from burst cells) were collected from the extracellular medium and transferred into Eppendorf tubes. Infected CHO cells were washed twice with PBS and the washing PBS was collected in the tubes containing the released parasites. Parasites were pelleted and resuspended in PBS and incubated with 50 nM SYTO9 (S34854) dye for 30 min at 37 °C with 5% CO_2_. Following this, the parasites were fixed using 4% PFA in PBS for 2 h at RT. A relative quantitation of the released parasite population was done via fluorescence-associated cell sorting (FACS) on a FACScan (BD Biosciences) equipped with a solid-state laser (Cytek, Cambridge, UK) at 485 nm excitation and 535 nm emission (FL1). Each plate also contained non-infected untreated controls (blanks: 0% released parasites), infected untreated controls (negative control: 100% released parasite) and reference controls (positive control: benznidazole at 20 µM). Results were expressed as % released parasites compared to the negative control. Each assay was carried out in three independent experiments each in triplicates.

### Biosafety

Experimental work with live *T. cruzi* were carried out following standard operating procedures in compliance with biosafety level 3* regulations (BSL3*) approved by the safety authorities of the University and Canton of Bern, Switzerland: The Standard Operational Procedures of the experiments were reported to the Swiss Authority Federal Office of Public Health (BAG).

### Comparison of EtOAc plant extract libraries

Hit rates were defined and calculated for both the Bolivian CD and the *DMM* extract libraries. For host cell cytotoxicity, the hit criterion was > 50% HeLa growth inhibition at 25 μg/mL (72 h incubation). The hit criterion for antichagasic activity was > 50% *T. cruzi* epimastigote growth inhibition at 25 μg/mL (72 h incubation). The criterion for potentially selective antichagasic activity was > 50% *T. cruzi* epimastigote growth inhibition at 25 μg/mL and < 50% inhibition of HeLa or CHO cellular proliferation at 25 μg/mL. The null hypothesis of the statistical test was that no relationship exists on the categorical variables (probability of hits) in the equally heterogeneous (genera and families) extract libraries, which were independent and with a reasonable sample size (>100 and <700). Results related to general cytotoxicity and selective antichagasic effects were compared with the Χ^2^ test and *P*-values were calculated. A *P*-value < 0.05 was considered as statistically significant. We investigated the phylogenetic distribution of plant species with the ggtree package (36) in the R environment Version 3.5.0 (http://www.R-project.org/).

### Microfractionation, bioactivity-guided isolation and structure elucidation

General Experimental Procedures. Optical rotations were measured in chloroform or methanol on a Jasco P-2000 digital polarimeter (Tokyo, Japan) with a 10 cm microcell. Nuclear magnetic resonance (NMR) spectra were recorded on a Bruker AVANCE III 500 MHz spectrometer (Billerica, CA, USA) operating at 500.13 MHz for 1H and 125.77 MHz for 13C. Measurements were performed with a 1 mm TXI probe at 18 °C. Data were processed with Bruker TopSpin 3.5 software. HR-ESIMS data were recorded in positive ion mode on a Thermo Scientific Orbitrap LQT XL mass spectrometer (Waltham, MA, USA). Centrifugal partition chromatography (CPC) was performed on an Armen Instrument (AlphaCrom, Rheinfelden, Switzerland) with coil volume 100 mL connected to a Varian pump model 210 (Agilent technologies, Santa Clara, CA, USA), a Varian 218 UV detector and a Varian fraction collector (model 704). Preparative reversed-phase high performance liquid chromatography (RP-HPLC) was carried out on a Puriflash 4100 system (Interchim, Montluçon, France) with a Waters (Milford, MA, USA) SunFire Prep C18 OBD column (5μm, 30 × 150mm i.d., guard column 10 × 20 mm i.d.). Semipreparative RP-HPLC was performed on an Agilent 1100 Series instrument with a DAD detector (Santa Clara, CA, USA) and a Waters (Milford, MA, USA) SunFire C18 column (5 μm, 150 × 10 mm i.d., guard column 10 × 10 mm i.d.). An HPLC system consisting of degasser, binary mixing pump, autosampler, column oven, and a diode array detector (all Shimadzu, Kyoto, Japan) connected via a T-splitter to an Alltech 3300 ELSD detector (Büchi, Flawil, Switzerland) and a Shimadzu 8030 triple quadrupole MS system with ESI and APCI interfaces was used for HPLC analysis. Data acquisition and processing were performed with LabSolution software (Kyoto, Japan). Separations were achieved with a SunFire C18 column (3.5 μm, 150 × 3.0 mm i.d., guard column 10 mm × 3.0 mm i.d.) (Waters, Milford, MA, USA). Microfractionation was carried out on the same instrument, but with a FC 204 fraction collector (Gilson, Middleton WI, USA) connected instead of the MS. Microfractions were collected into 96-deepwell plates. HPLC grade acetonitrile (Macron Fine Chemicals, Avantor Performance Materials, Phillipsburg, NJ, USA) and water from a Milli-Q water purification system (Merck, Darmstadt, Germany), and formic acid from Merck were used for HPLC separations. NMR spectra were recorded in DMSO-d6, methanol-d4 or chloroform-d (Armar Chemicals, Döttingen, Switzerland).

Microfractionation and isolation of natural products from selected extracts was carried out for subsequent testing on epimastigote viability/proliferation (72 h incubation assay) and in the parasite release assay. For microfractionation, extracts were dissolved in DMSO (10 mg/mL) and separated by gradient HPLC [0.1% aq. formic acid (A), 0.1% formic acid in acetonitrile (MeCN) (B); 0−30 min (5-100 % B); flow rate 0.4 mL/min; injection volume 4 x 30 μL; room temperature]. Optimized gradients were used for *Laurus nobilis* leaf extract [0−5 min (5-30 % B), 5−30 min (30-100 % B)] and *Sium sisarum* root extract [0−5 min (5-40 % B), 5−30 min (40-1000 % B)]. One-minute fractions were collected in 96-deep-well plates. Plates were dried in a Genevac EZ-2 evaporator (Ipswich, UK), and microfractions in wells redissolved in DMSO prior to bioassays. Compounds in the active time windows were isolated as follows: 200.6 mg of EtOAc extract of *Rumex crispus* rhizome were submitted to preparative RP-HPLC [water (A), MeCN (B); 0− 15 min (40−60 % B), 15−30 min (60 % B), 30-50 min (60-100 % B); flow rate 20 mL/min; injection volume 2 × 2 mL; room temperature] afforded compounds nepodin (28.2 mg, tR 19.3 min), torachrysone (2.7 mg, tR 23.9 min), emodin (1.67 mg, tR 24.7 min), and chrysophanol (8.0 mg, tR 38.9 min). 24.0 mg of EtOAc extract of S. sisarum roots were purified by semipreparative RP-HPLC [water (A), MeCN (B); 0−5 min (5-40 % B), 5−20 min (40-70 % B), 20-30 min (70 % B); flow rate 4 mL/min; injection volume 3 × 400 µL; room temperature] to yield falcarindiol (7.1 mg, tR 21.8 min). 30.0 mg of EtOAc extract of *Rheum rhaponticum* roots were separated by semipreparative RP-HPLC [water (A), MeCN (B); 0−20 min (5-60 % B), 20−30 min (60 % B); flow rate 4 mL/min; injection volume 3 × 500 µL; room temperature] to afford (6R, 7S)-costunolide (1.1 mg, tR 26.3 min). 2.0 g of *L. nobilis* leaf extract were separated by CPC [Hexane/EtOAc/MeOH/water, 7/3/5/5, v/v/v/v]; 0-60 min (descending mode), 60-72 min (ascending mode); flow rate 5 mL/min; rotation 2000 rpm; injection volume 10 mL]. Separation was monitored by the HPLC gradient mentioned above and detection at 210 nm. Active compounds were in fractions F2 (18-20 min), F6 (46-49 min), F9 (67-72 min). These fractions were further purified by semipreparative RP-HPLC [water (A), MeCN (B); flow rate 4 mL/min; injection volume 3 × 500 µL; room temperature]. F2 (32.0 mg) [0−45 min (25 % B)] yielded compounds (+)-reynosin (3.8 mg, tR 26.6 min), zaluzanin C (1.6 mg, tR 29.2 min), and santamarine (2.7 mg, tR 39.7 min). F6 (13.0 mg) [0−30 min (42 % B)] afforded (3S)-3-acetylzaluzanin C (4.2 mg, tR 23.8 min), and F9 (44.0 mg) [0−30 min (40-65 % B)] yielded costunolide (14.1 mg, tR 21.9 min), dehydrocostus lactone (5.3 mg, tR 23.7 min), and eremanthin (2.4 mg, tR 24.7 min).

*Nepodin*: amorphous solid; HR-ESI-MS m/z 217.0850 [M + H]^+^ (calcd for C_13_H_13_O_3_^+^, 217.0859).

*Torachrysone*: amorphous solid; HR-ESI-MS m/z 247.0963 [M + H]^+^ (calcd for C_14_H_15_O_4_^+^, 247.0965).

*Emodin*: amorphous solid; HR-ESI-MS m/z 271.0598 [M + H]^+^ (calcd for C_15_H_11_O_5_^+^, 271.0601).

*Chrysophanol*: amorphous solid; HR-ESI-MS m/z 255.0649 [M + H]^+^ (calcd for C_15_H_11_O_4_^+^, 255.0652).

*(3R, 8S)-Falcarindiol*: oil; [α]_D_^25^ +285 (0.1, CHCl_3_); APCI-MS m/z 262 [M + H]^+^.

*(6R, 7S)-Costunolide*: amorphous solid; [α]_D_^25^ +108 (0.1, CHCl_3_); HR-ESI-MS m/z 233.1527 [M + H]^+^ (calcd for C_15_H_21_O_2_^+^, 233.1536).

*(+)-Reynosin*: amorphous solid; [α]_D_^25^ +59 (0.2, CH_3_OH); ESI-MS m/z 249 [M + H]^+^.

*Zaluzanin C*: amorphous solid; [α]_D_^25^ +37 (0.1, CHCl_3_); ESI-MS m/z 247 [M + H]^+^.

*Santamarine*: amorphous solid; [α]_D_^25^ +83 (0.1, CH_3_OH); ESI-MS m/z 249 [M + H]^+^.

*(3S)-3-acetylzaluzanin C*: amorphous solid; [α]_D_^25^ +15 (0.1, CHCl_3_); ESI-MS m/z 289 [M + H]^+^.

*Dehydrocostus lactone*: amorphous solid; [α]_D_^25^ −6 (0.1, CHCl_3_); ESI-MS m/z 231 [M + H]^+^.

*Eremanthin*: amorphous solid; [α]_D_^25^ −276 (0.1, CHCl_3_); ESI-MS m/z 231 [M + H]^+^.

For the SAR analysis with 9,10-anthracenedione, the following commercial compounds were purchased (≥95% purity). Anthraquinone (**1**), alizarin (**4**), alizarin RedS (**5**), quinizarin (**6**), anthrarufin (**7**), anthraflavic acid (**9**), emodin (**2**), purpurin (**3**), rhein (**15**), diacerein (**16**), chrysophanol (**10**), disperse Red11 (**17**) and dantron (**8**) were all obtained from Sigma Aldrich, Switzerland. Aloe emodin (**11**), physcion (**12**), 2-hydroxy-3-methylanthraquinone (**13**) and 2-hydroxy-1-methylanthraquinone **(14)** were obtained from Toronto Research Chemical, ON, Canada. Aurantio obtusin (**18**) was obtained from AdooQ Biosciences LLC, CA, USA.

## Results

### Ethnopharmacological survey in Bolivia

Ethnomedical data were obtained from a total of 361 informants in three different geographic areas representing four ethnic groups (Fig. 1). In total, 152 research participants (5 Ayoreo, 68 Chiquitano, 54 Izoceño-Guaraní, 19 Quechua and 6 sellers from the city markets) reported ethnomedical knowledge related to the treatment of CD (Table 1). In our study, the gender of the informants was uniformly distributed, and we were not able to detect gender-specific knowledge related to the treatment of CD. When asked about CD and its symptoms, informants of all ethnic groups showed a lack of ethnomedical knowledge related to the pathophysiology of CD. Unlike other disease categories, such as dermatological affections or diseases of the digestive system, chronic CD was recognized unambiguously only after diagnosis by blood tests in the health centers or hospitals. Symptoms related to cardiomyopathies (general fatigue, shortness of breath, dizziness, chest pain, heart palpitations, high blood pressure and swelling of feet and legs) were generally associated to CD among the Chiquitano and Izoceño-Guaraní. Botanical drugs used for CD were most frequently associated with these symptoms. The survey resulted in more than 350 use reports for 79 plant taxa used in the context of CD (Table 2). Of these, 69 were identified to the species level, 9 plants were identified to the genus level, and one taxon was identified only to the family level. The recorded remedies comprised plant taxa distributed across 37 botanical families and 74 genera (Table 2). From the plants identified to the species level, 80% were native to Bolivia. Only 41 medicinal plant species (52%) were collected wild, while the others were obtained through cultivation or from local markets. Decoctions and infusions in cold and hot water, respectively, were the most common preparation forms for oral administration. Informants reported that the chronic symptoms were relieved after taking these plant remedies. The most cited plant species to be effective in the treatment of symptoms related to CD were *Senna chloroclada* (flowers, root) (Fig. 3) and *Tabebuia aurea* (bark), followed by the introduced and globally known medicinal plant species *Alpinia zerumbet* (rhizome) and *Cymbopogon citratus* (herb) (Table 2). Noteworthy, Chiquitano informants also reported a blend of several botanical drugs against CD, such as a combination of the barks of *Jacaranda cuspidifolia*, *T. aurea* and *Trichilia* sp. Particularly interesting in the context of this study was that few informants stated that certain botanical drugs could cure chronic CD, namely *S. chloroclada* (flowers, root), *Cochlospermum tetraporum* (bark, leaves) and *Bulnesia bonariensis* (bark). However, there was no consensus among independent informants. Even during prolonged stays, the use of these remedies was observed rarely by the authors. Only the use of *Galphimia brasiliensis* (root) (Fig. 3) could be repeatedly observed among the Chiquitano. To treat triatomine bites, topical application as poultices were mentioned (Table 2). Informants agreed that acute symptoms and inflammatory swellings (chagoma) at the bite sites occurred very rarely. Nocturnal triatomine bites were considered common and harmless. In highly affected communities (Izoceño-Guaraní and Ayoreo) informants stated that the bites were irritating but not painful and therefore remained mostly untreated. Only the taxa *Aloe vera*, *Croton* sp. and *Verbesina* sp were reported to be used for triatomine bites. *Bauhinia* sp., *S. chloroclada* and *Parthenium hysterophorus* were mentioned as a treatment of fever during first infections (Table 2). Besides plant based remedies, eleven Chiquitano informants reported the use of animal products, such as *Coragyps atratus* (black vulture; blood), *Tapirus* sp. (tapir; nails) and *Equus africanus asinus* (donkey; milk) to treat the chronic symptoms of CD. Few Izoceño-Guaraní informants also reported the use of grease of armadillos (*Dasypoda* spp.) against triatomine bites, and others applied alcohol topically. The plant taxa used for CD obtained in markets were common medicinal plants known to be used for a variety of diseases and ailments (Table 2).

**Figure 2.**
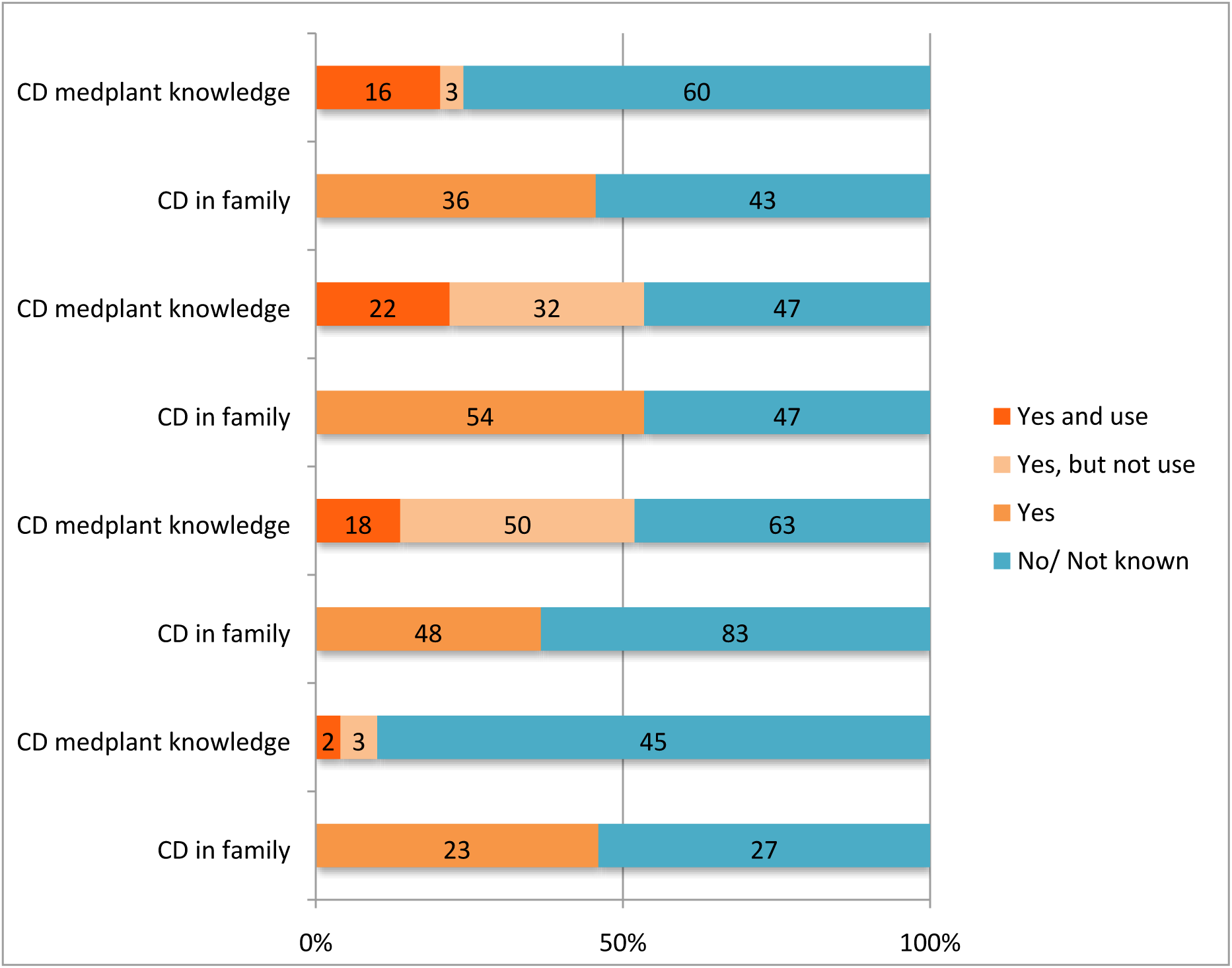
Graphical summary of the ethnomedical data. The number of informants from the four ethnic groups reporting knowledge and use of CD medicinal plants/agents (CD medplant knowledge), and the corresponding stated occurrence of CD in the family (CD in family) are shown.

**Figure 3.**
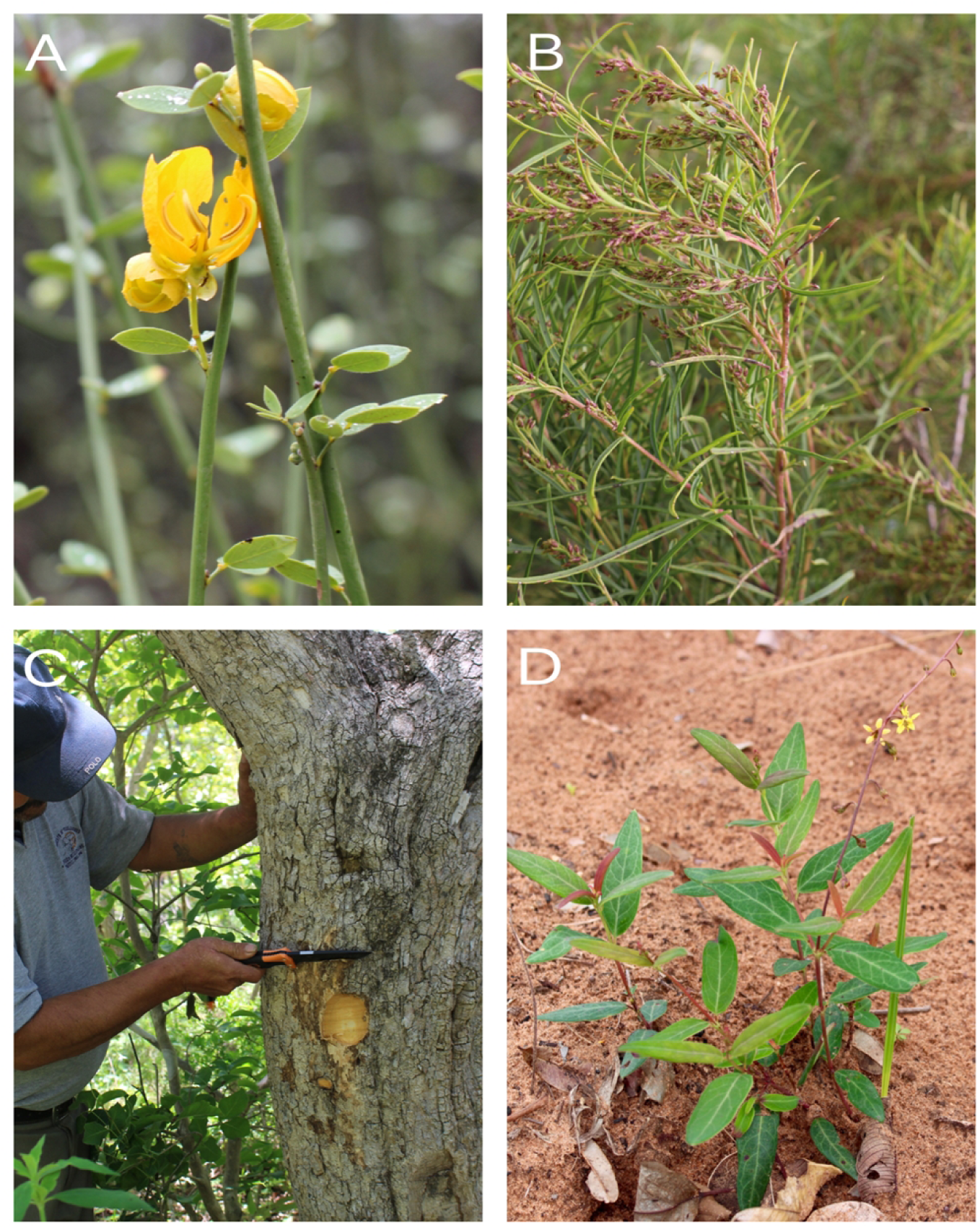
Selected Bolivian plant species used to treat symptoms related to CD. (A) *Senna chloroclada* flowers and roots were the most frequently cited botanical drugs among the Izoceño-Guaraní showing significant selective antichagasic effects on cellular parasite release *in vitro*. (B) *Acanthostyles buniifolius* was used by the Quechua and exhibited selective antichagasic activity *in vitro*. (C) Chiquitano informant cutting tree bark of *Jacaranda cuspidifolia* (D) The roots of *Galphimia brasiliensis* are used by the Chiquitano to treat symptoms related to CD.

**Table 1.** Socio-demographic characteristics of the informants in the surveyed rural areas reporting knowledge about CD medicinal plants*, number of use reports and reported taxa.

| Ethnicity | Geographical zone | Total | Gender |  | Age class |  |  | Use | Reported |
| --- | --- | --- | --- | --- | --- | --- | --- | --- | --- |
|  |  | Informants | F | M | < 40 | 41 to 60 | > 60 | reports | taxa |
| Ayoreo | Chiquitania | 5 | 3 | 2 | 1 | 4 | 0 | 5 | 3 |
| Chiquitano | Chiquitania | 68 | 33 | 35 | 22 | 24 | 22 | 199 | 38 |
| Guaraní | Chaco | 54 | 33 | 21 | 19 | 24 | 11 | 93 | 15 |
| Quechua | Inter-andean valleys | 19 | 10 | 9 | 8 | 9 | 2 | 42 | 31 |
| Total |  | 146 | 79 | 67 | 50 | 61 | 35 | 339 | 87 |

\* responses of women sellers in herbal markets in La Paz and Santa Cruz cities are not included in this table.

**Table 2.** List of plant species reported to be used in the treatment of Chagas disease and related symptoms by Ayoreo, Chiquitano, Guaraní and Quechua informants.

| Family<br>Genus/species | Vernacular name | Used<br>by | Part<br>used | No. use<br>reports | Application | Origin<br>(status) | Habit | Acute or<br>chronic<br>phase | Indication<br>Chagas<br>symptom | Voucher<br>specimen |
| --- | --- | --- | --- | --- | --- | --- | --- | --- | --- | --- |
| Amaranthaceae |  |  |  |  |  |  |  |  |  |  |
| *Dysphania ambrosioides* (L.) Mosyakin & Clemants | Caré, paico | C | L, R | 17 | Or | N (Cul) | herb | chro | CAR, GAS | ASMP33 |
| Anacardiaceae |  |  |  |  |  |  |  |  |  |  |
| *Astronium urundeuva* (Allemao) Engl. | Cuchi | C | B, L | 2 | Or | N (W) | tree | chro | CAR | ASMP56,<br>ASMP6 |
| *Schinus molle* L. | Molle | Q | L | 2 | Or, Tp | N (W) | tree | chro | CAR, GAS | ASMP90 |
| Annonaceae |  |  |  |  |  |  |  |  |  |  |
| *Annona nutans* (R. E. Fr.) R. E. Fr. | Sinini, Aratiku, Sorimimi | G | L | 5 | Or | N (W) | shrub | chro | CAR | ASMP11 |
| *Duguetia* sp. | Sinini | C | L | 12 | Or | n.d. (Cul) | tree | chro | CAR | ASMP41 |
| Apocynaceae |  |  |  |  |  |  |  |  |  |  |
| *Aspidosperma quebracho-blanco* Schitdl. | Cacha | C | B | 8 | Or | N (W) | tree | chro | CAR, FAT | ASMP35 |
| *Vallesia glabra* (Cav.) Link | Amarguillo, Arakuaembu | G | L | 3 | Or | N (W) | tree | chro | CAR | ASMP17 |
| Aristolochiaceae |  |  |  |  |  |  |  |  |  |  |
| *Aristolochia andina* F. Gonzáles & I. Vargas | Waje | Q | Br, L | 1 | Or | N (W) | vine | chro | CAR | ASMP94 |
| Asteraceae |  |  |  |  |  |  |  |  |  |  |
| *Acanthostyles buniifolius* (Hook. ex Hook. & Arn.) | Romero | Q | AP | 2 | Or | N (Cul) | shrub | chro | FAT | ASMP82 |
| *Achyrocline alata* (Kunth) DC. | Guacanqui, Wira wira | C, LP | AP | 6 | Or | N (Cul,Pur) | herb | chro | CAR | ASMP45,<br>ASMP103 |
| *Achyrocline hyperchlora* S. F. Blake | Wira wira | Q | AP | 2 | Or | N (W) | herb | chro | CAR | ASMP91 |
| *Ambrosia arborescens* Mill. | Altamisa | Q | L | 2 | Or | N (W) | subshru | chro | CAR | ASMP83 |
| *Baccharis genistelloides* (Lam.) Pers. | Carqueja, Kinsa k'ucho | LP, Q | AP | 3 | Or | N (Pur,W) | herb | chro | CAR, GAS | ASMP28,<br>ASMP85 |
| *Bidens andicola* Kunth | Misuka | Q | AP | 1 | Or | N (W) | herb | chro | PUR | ASMP89 |
| *Chromolaena connivens* (Rusby) R.M. King & H. Rob. | Sunchuj maman | Q | Fr, L | 1 | Or | N (W) | shrub | chro | FAT | ASMP75 |
| *Culcitium canescens* Humb. & Bonpl. | K'ia k'ia | LP | AP | 2 | Or | N (Pur) | herb | chro | CAR | ASMP102 |
| *Cynara* sp. | Alcachofa | LP, SC | AP | 2 | Or | n.d. (Pur) | herb | chro | CAR | ASMP13 |
| *Gochnatia boliviana* S. F. Blake | Melinco | Q | L | 1 | Or | N (W) | shrub | chro | CAR, FAT | ASMP87 |
| *Mutisia acuminata* Ruiz & Pav. | Chinchirkuma | Q | Br, L | 1 | Or | N (W) | shrub | chro | CAR | ASMP88 |
| *Parthenium hysterophorus* L. | Artemisa, Chupurujumo | C | AP | 1 | Or | N (W) | herb | acu, chro | FEV, PUR | ASMP59 |
| *Pluchea sagittalis* (Lam.) Cabrera | 4 Cantos | C | Fl, L | 1 | Or | N (Cul) | shrub | chro | CAR | ASMP36 |
| *Schkuhria pinnata* (Lam.) Kunza ex Thell. | Jayaj pichana | Q | AP | 2 | Or | N (W) | herb | chro | CAR, FAT | ASMP76 |
| *Sonchus oleraceus* L. | Diente de Leon | C, LP | AP | 3 | Or | I (Pur) | herb | chro | CAR | AMP12,<br>ASMP30 |
| *Verbesina* sp. | Tabaco | Q | L | 1 | Tp | n.d. (W) | tree | acu | BIT | ASMP93 |
| **Bignoniaceae** |  |  |  |  |  |  |  |  |  |  |
| *Handroanthus impetiginosus* (Mart. ex DC.) Mattos | Tajibo, Tajibo negro | C, G | B, Fl | 3 | Or | N (W) | tree | chro | CAR | ASMP18, |
| *Jacaranda cuspidifolia* Mart. | Paraparau | C | B | 6 | Or | N (W) | tree | chro | CAR, FAT | ASMP34 |
| *Tabebuia aurea* (Silva Manso) Benth. & Hook. F. ex S. | Paratodo, Alcornoque | C, A | B | 30 | Or | N (Cul) | tree | chro | CAR, FAT, | ASMP9 |
| **Bixaceae** |  |  |  |  |  |  |  |  |  |  |
| *Bixa orellana* L. | Urugu | C, G | L | 4 | Or | N (W) | tree | chro | CAR | ASMP26 |
| **Caricaceae** |  |  |  |  |  |  |  |  |  |  |
| *Carica papaya* L. | Papaya | C, G | Fl | 3 | Or | I (Cul) | tree | chro | CAR, GAS | ASMP38 |
| **Cochlospermaceae** |  |  |  |  |  |  |  |  |  |  |
| *Cochlospermum tetraporum* Hallier f. | Kuari, Pela pela | G | B, L | 8 | Or | N (Cul) | tree | chro | CAR | ASMP22 |
| **Cucurbitaceae** |  |  |  |  |  |  |  |  |  |  |
| *Momordica charantia* L. | Balsamina | C | AP | 3 | Or | I (Cul) | vine | chro | CAR | ASMP58 |
| **Ephedraceae** |  |  |  |  |  |  |  |  |  |  |
| *Ephedra americana* Humb. & Bonpl. Ex Willd. | Pisqo simi | Q | AP | 1 | Or | N (W) | shrub | chro | FAT | ASMP96 |
| **Euphorbiaceae** |  |  |  |  |  |  |  |  |  |  |
| *Croton andinus* Muell. Arg. | Taporita, Tupeicha | G | R | 5 | Or | N (W) | herb | chro | CAR | ASMP21 |
| *Croton* sp. | K'uru k'uru | Q | La | 1 | Tp | n.d. (W) | herb | acu | BIT | ASMP92 |
| *Euphorbia serpens* Kunth | Quebra Pedra | C | Wh | 4 | Or | N (Cul) | herb | chro | CAR, FAT | ASMP49 |
| *Jatropha curcas* L. | Piñón | C | L | 1 | Or | I (Cul) | treelet | chro | FAT | ASMP39 |
| **Fabaceae** |  |  |  |  |  |  |  |  |  |  |
| *Acacia aroma* Gillies ex Hook. & Arn. | Tusca | G | B, Fl | 2 | Or | N (Cul) | tree | chro | CAR | ASMP24 |
| *Acacia* sp. | Karikari | C | B | 1 | Or | n.d. (Cul) | tree | chro | PUR | ASMP43 |
| *Bauhinia* sp. | Patecabra | C | L | 1 | Or | n.d. (Cul) | shrub | acu, chro | FEV, PUR | ASMP42 |
| *Copaifera langsdorffii* Desf. | Copaibo | C, SC | B | 5 | Or | N (W) | tree | chro | CAR | ASMP51 |
| *Crotalaria incana* L. | Amorocita | Q | L | 1 | Or | N (W) | herb | chro | FAT | ASMP95 |
| *Hymenaea courbaril* L. | Paquíó | C | B | 3 | Or | N (W) | tree | chro | CAR, FAT | ASMP55 |
| *Pterodon* sp. | Pezoe | C | Se | 2 | Or | N (W) | tree | chro | FAT | ASMP52 |
| *Senna chloroclada* (Harms) H. S. Irwin & Barneby | Lanza lanza, Mbuijare, Retama | G | Fl, R, | 45 | Or | N (W) | shrub | acu, chro | CAR, FEV | ASMP10 |
| *Spartium junceum* L. | Retama | A, SC | AP | 3 | Or | I (Pur) | shrub | chro | CAR | ASMP100 |
| **Gesneriaceae** |  |  |  |  |  |  |  |  |  |  |
| *Gloxinia gymnostoma* Griseb. | Ortelón | C | AP | 2 | Or | N (Cul) | herb | chro | CAR, FAT | ASMP40 |
| **Gramineae** |  |  |  |  |  |  |  |  |  |  |
| *Cymbopogon citratus* (DC.) Stapf | Paja de cedrón | C, G, SC | L | 25 | Or | I (Cul) | herb | chro | CAR, FAT | ASMP31 |
| **Labiatae** |  |  |  |  |  |  |  |  |  |  |
| *Clinopodium axillare* (Rusby) Harley | Huayra Muña | Q | Br, Fl, L | 2 | Or | N (Cul) | subshru | chro | CAR | ASMP80 |
| *Lepechinia vesiculosa* (Benth.) Epling | Raq'acho | Q | L | 1 | Or | N (W) | shrub | chro | CAR | ASMP81 |
| *Minthostachys ovata* (Briq.) Epl. | Muña | Q | AP | 3 | Or | N (Cul) | subshrub | chro | FAT, GAS | ASMP69 |
| *Ocimum americanum* L. | Albahaca | C | Wh | 2 | Or | I (Cul) | herb | chro | CAR | ASMP48 |
| *Aloe vera* (L.) Burm.f. | Sábila, karaguata guasu | C, G | La | 5 | Tp | I (Cul) | herb | acu | BIT, PUR | ASMP25 |
| Linaceae |  |  |  |  |  |  |  |  |  |  |
| *Linum usitatissimum* L. | Linaza | Q | Fr | 2 | Or | I (Cul) | herb | chro | PUR | ASMP71 |
| Malpighiaceae |  |  |  |  |  |  |  |  |  |  |
| *Galphimia brasiliensis* (L.) A. Juss. | Masiaré | A, C | R | 11 | Or | N (Cul) | herb | chro | CAR, FAT | ASMP32 |
| sterile | Azucaró | C | B | 4 | Or | n.d. (W) | tree | chro | CAR | ASMP54 |
| Malvaceae |  |  |  |  |  |  |  |  |  |  |
| *Malva parviflora* L. | Malva | LP, Q | AP | 2 | Or | I (Cul,Pur) | herb | chro | CAR, FAT, | ASMP77, |
| Meliaceae |  |  |  |  |  |  |  |  |  |  |
| *Trichilia* sp. | Tipa | C, SC | B | 11 | Or | n.d. (Cul) | tree | chro | CAR | ASMP53 |
| Myrtaceae |  |  |  |  |  |  |  |  |  |  |
| *Myrcianthes callicoma* McVaugh | Guapurú | Q | L | 1 | Tp | N (W) | tree | chro | FAT | ASMP73 |
| *Myrcianthes pseudomato* (D. Legrand) McVaugh | K'arasacha | Q | L | 2 | Or | N (W) | tree | chro | CAR, GAS | ASMP72 |
| *Plinia cauliflora* (Mart.) Kausel | Guapurú | C | L | 1 | Or | N (Cul) | tree | chro | CAR | ASMP60 |
| Oxalidaceae |  |  |  |  |  |  |  |  |  |  |
| *Hypseocharis pimpinellifolia* Remy | Sultaki | Q | R | 1 | Tp | N (W) | herb | chro | FAT | ASMP70 |
| Papaveraceae |  |  |  |  |  |  |  |  |  |  |
| *Argemone subfusiformis* G. B. Ownbey | Cardo Santo | C, G | Fl | 3 |  | N (W) | herb | chro | CAR | ASMP20 |
| *Bocconia integrifolia* Bonpl. | Turumi | Q | L | 1 | Or | N (W) | tree | chro | FAT | ASMP97 |
| Passifloraceae |  |  |  |  |  |  |  |  |  |  |
| *Passiflora cincinnata* Mast. | Pachío, Mburukuya | C, G | Fl, L, R | 13 | Or | N (W) | vine | chro | CAR, FAT | ASMP16 |
| Piperaceae |  |  |  |  |  |  |  |  |  |  |
| *Piper rusbyi* C. DC. | Matico | C | L | 7 | Or | N (Cul,W) | shrub | chro | FAT, PUR | ASMP50, |
| Plantaginaceae |  |  |  |  |  |  |  |  |  |  |
| *Plantago major* L. | Llantén | C | Fl, L | 3 | Or | I (Cul) | herb | chro | CAR | ASMP47 |
| Polygalaceae |  |  |  |  |  |  |  |  |  |  |
| *Monnina wrightii* A. Gray | T'ian t'ian | Q | L | 1 | Or | N (W) | herb | chro | CAR | ASMP84 |
| Rosaceae |  |  |  |  |  |  |  |  |  |  |
| *Rubus* sp. | Kari kari | LP | AP | 2 | Or | n.d. (Pur) | herb | chro | CAR | ASMP101 |
| Rutaceae |  |  |  |  |  |  |  |  |  |  |
| *Heterophyllaea lycioides* (Rusby) Sandwith | Kapi | Q | L | 1 | Or | N (W) | shrub | chro | GAS | ASMP86 |
| *Ruta chalepensis* L. | Ruda | Q | L | 1 | Or | I (Cul) | herb | chro | GAS, PUR | ASMP99 |
| *Zanthoxylum coco* Gillies ex Hook. F. & Arn. | Chirimolle | Q | L | 1 | Or | N (W) | tree | chro | CAR | ASMP74 |
| Solanaceae |  |  |  |  |  |  |  |  |  |  |
| *Cestrum parqui* L'Her. | Andrés Huaylla | LP, Q | AP, L | 2 | Or | N (Cul,Pur) | shrub | chro | CAR, GAS | ASMP78, |
| *Solanum palinacanthum* Dunal | Pica Pica | C | L, R | 1 | Or | N (Cul) | subshru | chro | PUR | ASMP44 |
| Urticaceae |  |  |  |  |  |  |  |  |  |  |
| *Urtica urens* L. | Ortiga | LP | AP | 1 | Or | I (Pur) | herb | chro | CAR | ASMP29 |
| Valerianaceae |  |  |  |  |  |  |  |  |  |  |
| *Valeriana potopensis* Briq. | Jama jama | Q | B, L | 1 | Or | N (W) | herb | chro | CAR | ASMP98 |
| Verbenaceae |  |  |  |  |  |  |  |  |  |  |
| *Aloysia citriodora* Palau | Cedrón | C, Q | L | 4 | Or | N (Cul) | shrub | chro | CAR | ASMP79 |
| *Aloysia polystachya* (Griseb.) Moldenke | Poleo | C | AP | 2 | Or | N (Cul) | shrub | chro | CAR | ASMP37 |
| Zingiberaceae |  |  |  |  |  |  |  |  |  |  |
| *Alpinia zerumbet* (Pers.) B. L. Burtt & R. M. Sm. | Colonia | C, G, SC | Fl | 26 | Or | I (Cul) | herb | chro | CAR | ASMP19 |
| Zygophyllaceae |  |  |  |  |  |  |  |  |  |  |
| *Bulnesia bonariensis* Griseb | Guayacán, Guayacán Morado | C, G | B | 3 | Or | N (Cul) | tree | chro | CAR | ASMP23 |

Abbreviations: A:Ayoreo; C:Chiquitano; G:Guaraní; LP:La Paz; SC:Santa Cruz; AP:aerial parts; B:bark; Br:branches; Fl:flowers; Fr:fruit; La:latex; L:leaves; R:roots; Se:seeds; Wh:whole plant; Or:oral application; Tp:topical application; N:native; I: introduced; n.d.: not determined; Cul:cultivated; Pur:purchased at market; W:wild; chro:chronic; acu:acute; BIT:Vinchuca bite; CAR:cardiovascular symptoms; FEV:fever; GAS:gastrodigestive symptoms; FAT:fatigue; PUR:purifier/blood purifier/to strengthen the blood

The Quechua and Ayoreo reported relatively fewer remedies to manage CD as compared to the Izoceño-Guaraní and Chiquitano (Table 1), despite the high prevalence of CD in their communities (> 40%; Fig. 2). The vast majority of the Ayoreo interviewed in this study did not report any botanical drugs used in the context of CD (Fig. 2). The visited Ayoreo communities were evangelic or catholic Christians that stopped practicing shamanism decades ago. Generally, they had no or little knowledge about traditional medicinal agents. Although Quechua informants have been educated on the association between triatomine bugs and CD by institutional anti-CD campaigns (41), they had a very limited understanding about the transmission mechanisms and symptoms of the illness. There was poor consensus among Quechua informants and we obtained only one or two use-reports for each plant taxon, with the exception of *Minthostachys ovata* aerial parts (3 use-reports, Table 2). Most of the reported plants used by the Quechua were native to the region (86% of the total number of identified taxa at species level) and were gathered from the wild (74%). Among the Quechua, traditional healers (curanderos) generally carried out the preparation of botanical drugs. Although there were specialized healers among the Izoceño-Guaraní (paye) and Chiquitano communities, the knowledge and use of botanical drugs was not restricted to them. Guaraní and Chiquitano community members used botanical drugs to alleviate CD-associated symptoms because they lacked alternatives, and ethnomedical remedies were easily accessible. Especially in the Guaraní communities there was a general lack of primary health care, and travelling to Santa Cruz posed significant difficulties for patients.

### Analysis of plant libraries

#### Biological profiling of CD botanical drug library, collected in Bolivia

Crude EtOAc extracts obtained from 115 botanical drugs, representing 79 plant species and collected on the basis of their use in the management of CD (Table 2), were dissolved in DMSO and tested *in vitro* for their antitrypanosomal activity against *T. cruzi* epimastigotes and procyclic *T. b. brucei*, as well as for general cytotoxic effects in HeLa and Raw264.7 cells. Antiproliferative IC_50_ values < 25 μg/mL were considered biologically significant. To define selective hits, a cutoff of 50% inhibition (HeLa antiproliferative effect) at 25 μg/mL was used. IC_50_ values were determined only for selective hits. Supplementary Table 1S summarizes the results obtained in the *in vitro* bioassays with the plant extracts. Phylogenetic distribution of reported anti-CD plant taxa and screening results are shown in Fig. 4. In general, the *T. b. brucei* strain was very sensitive towards the tested extracts and, rather surprisingly, 90 extracts (78%) exhibited antitrypanosomal activity against procyclic *T. b. brucei* (Fig. 4). In contrast, only 20 (17%) extracts inhibited *T. cruzi* epimastigotes with IC_50_ values ≤25 µg/mL. Seven extracts (6%) showed pronounced (IC_50_ < 10 μg/mL) and 13 extracts (10%) good (IC_50_ ≤ 25μg/mL) antitrypanosomal activity (Fig. 4). The most potent extracts were those of *Cynara* sp. aerial parts (2 μg/mL), *Acanthostyles buniifolius* aerial parts (2 μg/mL) and *Gochnatia boliviana* leaves (4 μg/mL) (Fig. 4). The extracts from roots and flowers of *S. chloroclada*, the most frequently cited anti-CD botanical drugs among the Guaraní, were not active against epimastigotes (IC_50_s > 50 μg/mL), while an extract of the aerial parts of this plant was moderately active. The *S. chloroclada* flower extract showed significant inhibitory effects on parasite release in the infection assay (15 μg/mL were equally effective as 20 μM of benznidazole) without being cytotoxic to host cells (Suppl. Table S1). The ESI-MS scan and a positive TLC Borntrager reaction indicated that this plant contained anthraquinones and possibly anthrones and its glycosides. Since many extracts exhibited significant general cytotoxicity (Fig. 4), the observed toxicity against epimastigotes could also be due to non-specific effects. Moreover, cytotoxic extracts (27.8%) could not be analyzed in the trypomastigote infection assay. Particularly cytotoxic extracts were from *Ambrosia arborescens* leaves, *G. boliviana* leaves, *G. boliviana* branches, *Bocconia integrifolia* leaves, *B. integrifolia* roots, *Cynara* sp. aerial parts, and *P. hysterophorus* aerial parts. The only highly active and selective antitrypanosomal extracts were from *Pterodon* sp. seeds, *Sonchus oleraceus* leaves, *A. buniifolius* aerial parts, *Aloysia polystachya* aerial parts and *Gloxinia gymnostoma* aerial parts (Fig. 4). The extracts of these taxa showed also significant inhibitory effects in the *T. cruzi* infection assay (Suppl. Table S1).

**Figure 4.**
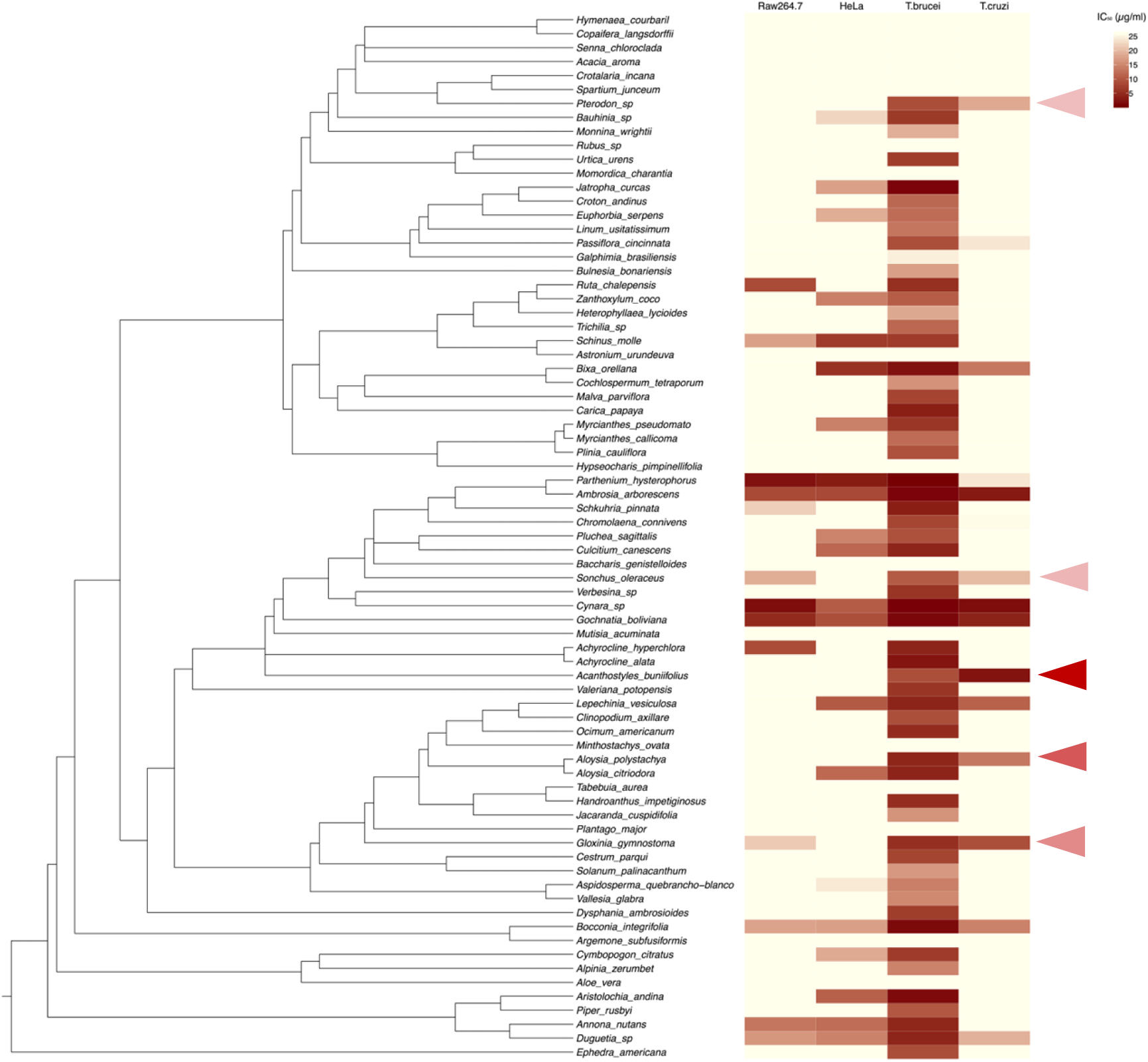
Phylobioactivity profiling of the EtOAc extract library generated from Bolivian medicinal plants used to treat symptoms of CD (reported plant part used). Few extracts showed selective toxicity towards *T. cruzi* epimastigotes with IC_50_ values below 20 µg/mL (arrows). There was a high of extracts toxic for procyclic *T. brucei*, and many showed antiproliferative effects in HeLa and Raw264.7 cells. The Leguminosae was the only subfamily cluster (top) showing no bioactivities up to 25 µg/mL. In the sesquiterpene lactone rich family Asteraceae, only *Acanthostyles buniifolius* showed selective antitrypanosomal effects. Data represent profiling values from at least two independent screening assays each performed in triplicates.

As initial IC_50_ values were obtained in anti-proliferation assays with *T. cruzi* epimastigotes, we next screened the extracts lacking general cytotoxicity in the cellular infection assay. In order to validate the antitrypanosomal activity on the mammalian stage forms that are relevant for the disease, extracts were tested for their ability to inhibit *T. cruzi* parasite release assay in CHO cells. Amastigotes develop intracellularly, differentiate into trypomastigotes and leave the host cell. Here we quantified the amount of trypomastigotes released (and residual amastigotes in case cells burst prematurely). To that aim, we developed a versatile FACS-based assay by staining released parasites in supernatant using a DNA stain. Benznidazole was used to validate the assay system (Fig. 5). Although the assay showed significant variability (± 30%), due to the nature of independent *T. cruzi* infections and overall variability in replication efficiencies, a reliable IC_50_ value for benznidazole could be obtained (2 ± 0.4 µM), which was in agreement with previously published data (58). Consequently, we employed this assay for our profiling of the CD botanical drug library, whereby initial testing of the extracts was performed at a single concentration (15 μg/mL). As shown in Table 1S, extracts which were active against *T. cruzi* epimastigotes were generally also active in the infection assay, with the exception of *S. chloroclada*. The pre-screening with epimastigotes was thus considered suitable for the discovery of antichagasic compounds able to inhibit different stages of the infection/replication cycle.

**Figure 5.**
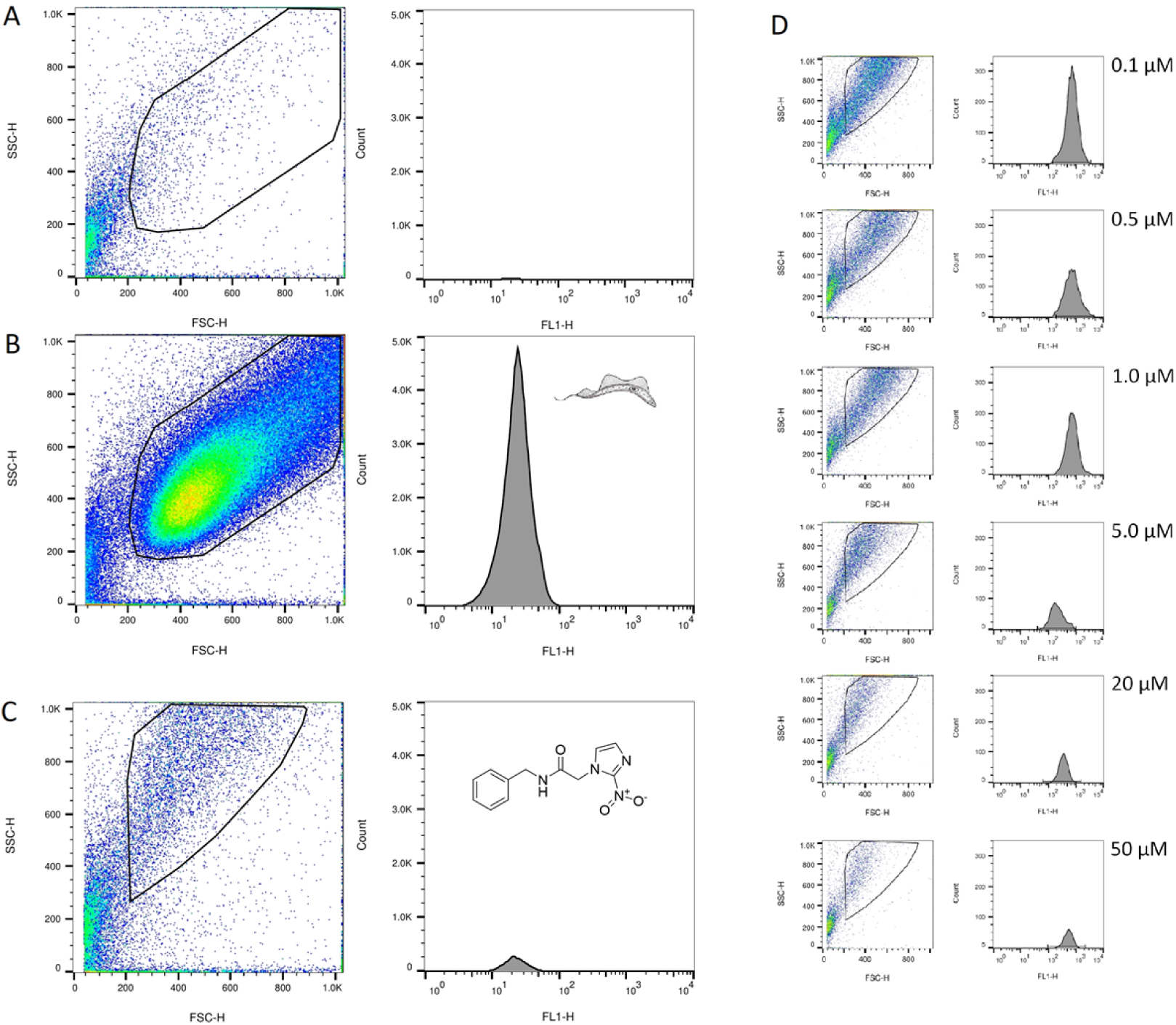
FACS-based trypomastigote infection assay in CHO cells. CHO cells were infected with trypomastigotes (10:1) for 24 h, washed, and incubated in low FBS (1%) medium for 6 days. Extracts dissolved in DMSO, vehicle and positive controls were added after washing and left for 6 days. Parasites released into the media at 6dpi were stained with SYTO™ green fluorescent nucleic acid stain, fixed with 4% paraformaldehyde and collected for FACS analysis. (A) Negative control of non-infected cells at 6 dpi showing FSC/SSC and FL1 histograms. (B) Positive control showing fluorescence associated to parasites at 6 dpi showing FSC/SSC and FLI1 histograms. (C) Representative example of benznidazole (Bnz) treatment (20 µM) at 6 dpi showing FSC/SSC and FL1 histograms illustrative of reduced parasite release. (D) Illustration of concentration-dependent effect by Benznidazol (Bnz) leading to decrease of parasites in medium at 6 dpi with a calculated IC_50_ value of 2 ± 0.4 µM (maximal efficacy about 75%).

#### Comparative profiling of the DMM library from the Mediterranean

To validate the ethno-directed approach for bioprospecting antichagasic activity, we compared the results obtained for botanical drugs used in a CD context with those of 660 botanical drugs described in Dioscorides’ *DMM*. Importantly, the botanical drugs mentioned in *DMM* show no traditional association with CD. EtOAc extracts were tested at 25 μg/mL *in vitro* against *T. cruzi* epimastigotes, and on HeLa cells for cytotoxicity (Fig. 6, Suppl. Table 2S). The extracts were considered active (i.e. hits) when the percentage of inhibition was ≥ 50% at 25 μg/mL. A total of 59 (8.9%) extracts exhibited antitrypanosomal activity while 102 (15.5%) extracts were cytotoxic towards HeLa cells (Fig. 6; Suppl. Table 2S). Among the antitrypanosomal hits, only 27 extracts (4.1%) were selectively toxic for *T. cruzi* epimastigotes. Extracts with apparent selectivity were those obtained from *Levisticum officinale* fruits, *Opopanax chironium* roots, *Glebionis coronaria* flowers, *Tanacetum parthenium* flowers, *Convolvulus scammonia* roots, *Iris foetidissima* seeds, *Laurus nobilis* fruits, *Rheum rhaponticum* roots, *Rumex crispus* roots and *Ruta chalepensis* roots. Extracts with selective toxicity for *T. cruzi* epimastigotes were considered potential hits and their antichagasic activity was subsequently determined at 15 μg/mL in the trypomastigote infection assay in CHO cells. All extracts with the exception of those obtained from *A. vera* resin, *L. nobilis* leaves and *L. albus* roots were also active in the infection assay (> 50 % inhibition of parasite release at 15 μg/mL).

**Figure 6.**
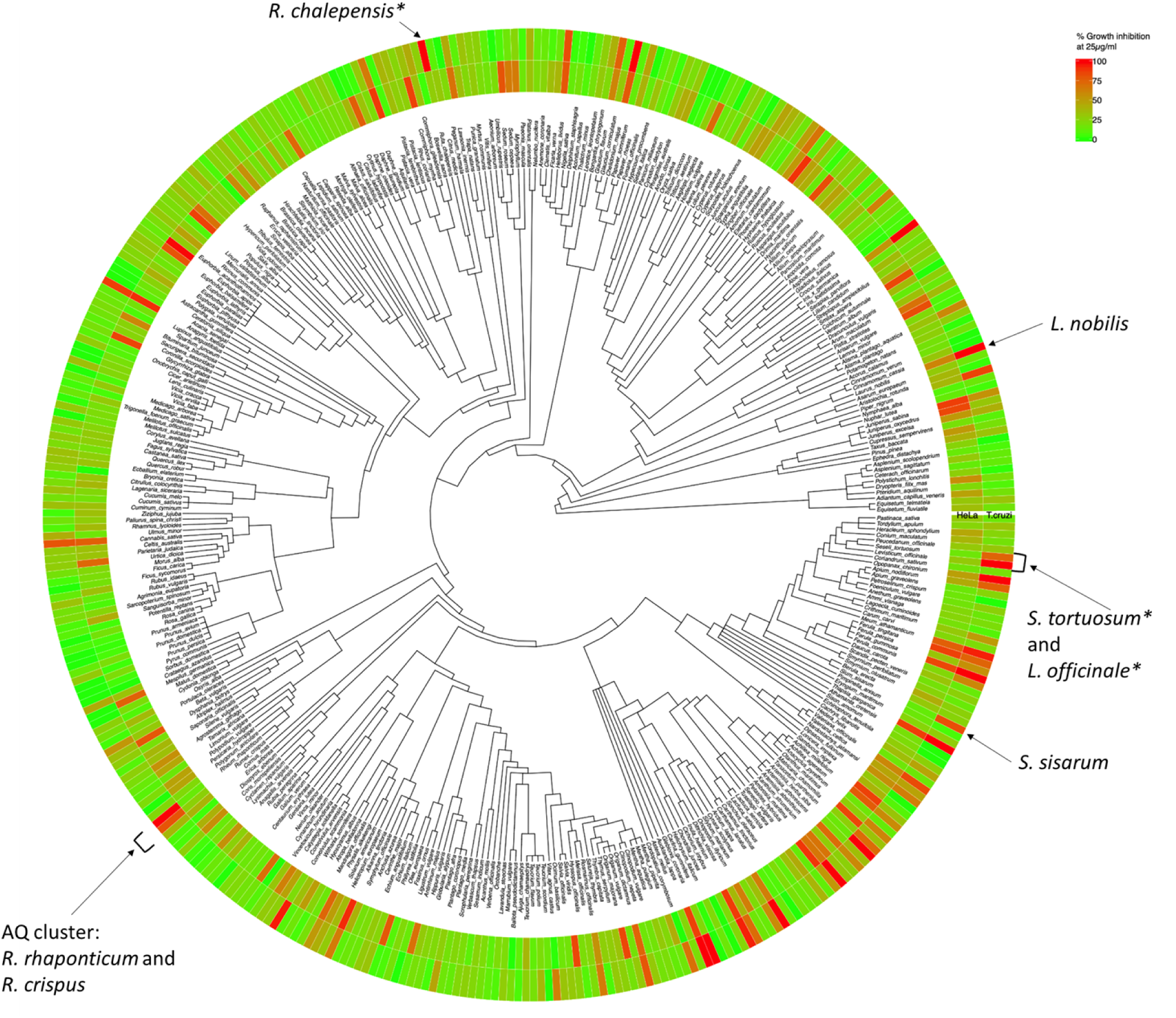
Phylobioactivity tree displaying phylogenetic relationships associated with bioactivities. The outer ring shows growth inhibition on *T. cruzi* epimastigotes, and the inner ring shows grows inhibition of HeLa cells (both at 25 µg/mL; most active plant part shown only). A hypothetical coumarin cluster (*) (*Ruta chalepensis root*, *Levisticum officinale seeds* and *Seseli tortuosum* root) and the anthraquinone cluster (*Rhumex crispus* and *Rheum rhaponticum* rhizoma) are visible. *Laurus nobilis* root and fruits and *Sium sisarum* root (microfractionated) are indicated in the phylogenic tree. Detailed data on plant species and activities are shown in supplementary Table 2S.

#### Comparison of extract libraries derived from different ethnopharmacological contexts

In an attempt to pharmacologically validate the ethno-directed approach, hit rates were calculated for both the CD and *DMM* botanical drug libraries. The Pearson Χ^2^ test was applied for statistics. Results show that there is a significantly higher probability (17.4% vs. 8.9%, *P* = 0.0057) of detecting antichagasic (*T. cruzi* epimastigotes) extracts when the plant had a reported use against CD. However, we also found a significantly higher percentage of cytotoxic (IC_50_ ≤ 25 μg/mL) extracts among CD botanical drugs (27.8% vs. 15.5%, *P* = 0.0012). Taking the importance of selectivity into account, the hit rate of selective antichagasic extracts *in vitro* was not considered statistically different (*P* = 0.079) between the Bolivian CD (7.8%) and the *DMM* (4.1%) extract libraries. The two libraries shared 20 genera and included the identical species *S. oleraceus*, *Spartium junceum*, *A. vera*, *Linum usitatissimum* and *R. chalepensis* (Suppl. Tables 1S and 2S). Only *S. oleraceus* and *S. junceum* aerial parts were also identical botanical drugs.

#### Microfractionation of selected extracts and identification of antitrypanosomal natural products

For the isolation of potentially antichagasic metabolites related to the *DMM* library, active extracts from different phylogenetic clusters were selected (Fig. 6). A major phylogenetic hotspot showing trypanocidal selectivity was the ‘anthraquinone cluster’ with *R. crispus* (curly dock) and *R. rhaponticum* (rhapontic rhubarb) rhizomes from the Polygonaceae family (Fig. 6). As anticipated, anthraquinones and naphthoquinones were major active principles in *R. crispus* (Fig. 7) and *R. rhaponticum* (not shown). The latter also yielded the SL (6*R*, 7*S*)-costunolide. *L. nobilis* (bay) leaves from the Lauraceae family were also studied. Noteworthy, all plant parts of *L. nobilis* showed significant antichagasic effects (Suppl. Table S2). The *L. nobilis* leaf extract, which inhibited epimastigote proliferation but was inactive in the infection assay, yielded the known SLs (+)-reynosin, zaluzanin C, santamarine, (3*S*)-3-acetylzaluzanin C, dehydrocostus lactone and eremanthin. Since all of these SLs showed antiproliferative effects in CHO cells (Table 3) we tested them in the infection assay at the subcytotoxic concentration of 5 µM only (Table 3). *Sium sisarum* root (skirret) from the Apiaceae family showed overall good antichagasic selectivity. However, phytochemical annotation in this case was more complex, since the polyacetylene (3*R*, 8*S*)-falcarindiol, a major metabolite from *S. sisarum*, was isolated but inactive in all assays. As illustrated with *R. crispus*, the most potent antichagasic principle was the anthraquinone emodin which completely inhibited *T. cruzi* epimastigote proliferation during the bioactivity-guided isolation and also inhibited parasite release (Fig. 7 and Table 3). Nepodin from *R. crispus* inhibited epimastigote proliferation (IC_50_ = 28.7 ± 13.3 µM) but was not equally efficacious in inhibiting the parasite release in the infection assay (∼30% inhibition at 5 µM).

**Figure 7.**
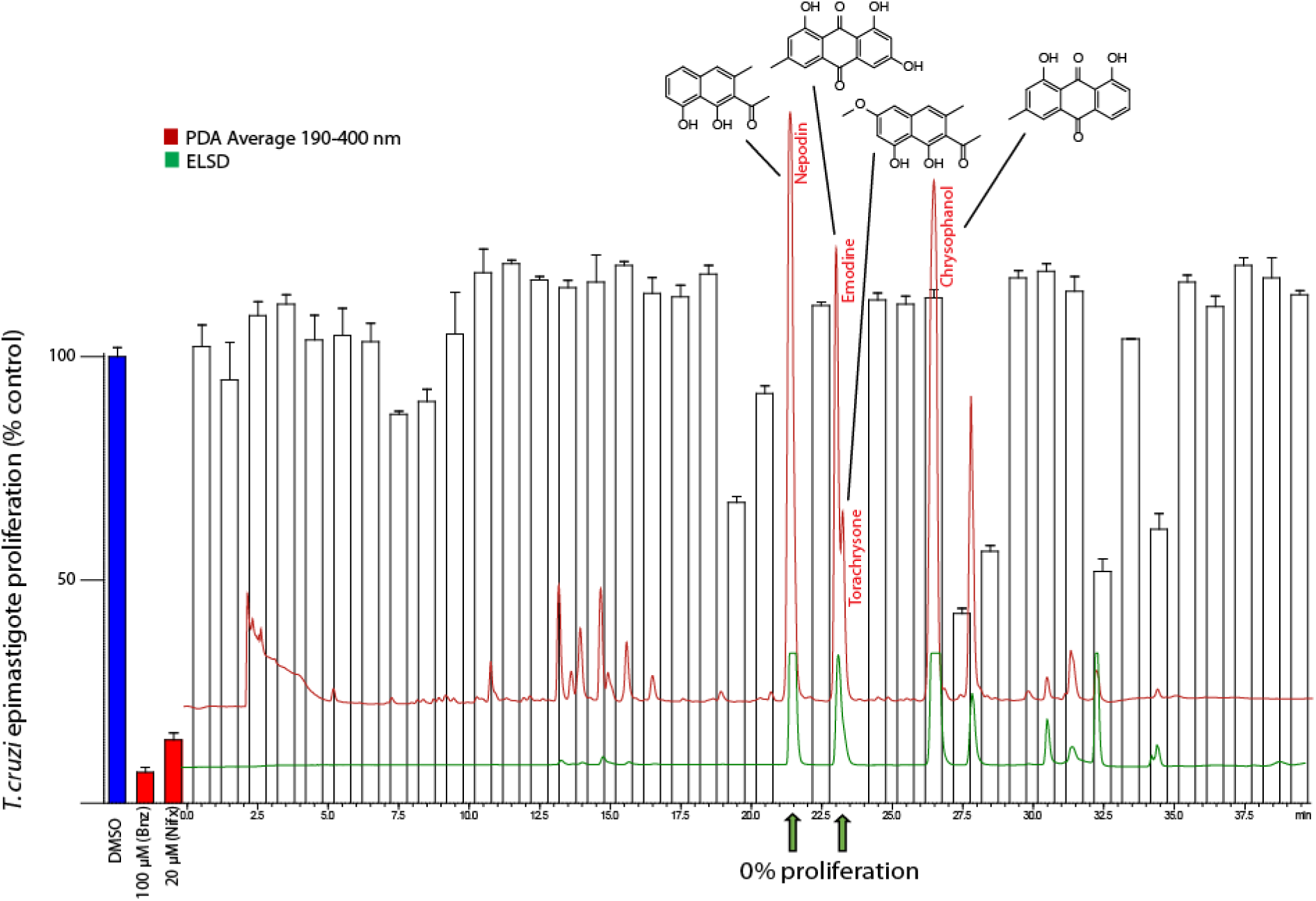
Bioactivity-guided microfractionation exemplified by *R. crispus* using photo diode array (PDA) and evaporative light scattering detectors (ELSD). Isolation of antichagasic metabolites was based on epimastigote proliferation inhibition. A limitation of this qualitative approach are false negatives (shown here with chrysophanol) due to low concentrations. Nepodin and emodin/torachrysone were identified and isolated from fully active fractions (0% cell viability), the moderately active chrysophanol was identified in a negative fraction.

**Table 3.** *In vitro* antiproliferative activity of compounds isolated from plant extracts on *T. cruzi* epimastigote-stage (72 h) and trypomastigotes release (6 dpi). Cytotoxic (antiproliferative) effects of the compounds were assessed on CHO host cells after 72 h. CHO cells cultured in medium without FBS showed a reduced cytotoxicity towards sesquiterpene lactones than proliferating CHO cells. Data shown are mean values ± SD of at least three independent experiments each performed in triplicates.

| Isolated cpd | IC<sub>50</sub> epimastigotes<br>[μM] | % inhibition of parasite<br>release<br>at 5 μM | IC<sub>50</sub> CHO cells<br>[μM] | IC<sub>50</sub> CHO cells<br>without FBS<br>[μM] |
| --- | --- | --- | --- | --- |
| Nepodin * | 28.7 ± 13.3 | 34.7 ± 30.9 | >100 | n.d |
| Torachryson * | >50 | 25.9 ± 15.5 | >100 | n.d |
| Emodin * | 14.1 ± 8.2 | 61.5 ± 18.2 | >100 | n.d |
| Falcarindiol <sup>†</sup> | >50 | 0 | >100 | n.d |
| Costunolide <sup>‡</sup> | 7.4 ± 5.9 | 0 | 10.9 ± 3.7 | 35.4 ± 13.8 |
| Reynosin <sup>#</sup> | >50 | 50.0 ± 47.1 | 31.3 ± 8.0 | 66.1 ± 1.2 |
| Santamarine <sup>#</sup> | 19.5 ± 9.3 | 55.9 ± 46.7 | 13.2 ± 4.3 | 39.4 ± 1.5 |
| Zaluzanin C <sup>#</sup> | 6.7 ± 0.7 | 71.0 ± 7.5 | 6.6 ± 1.7 | 19.2 ± 5.2 |
| 3-acetylzaluzanin C <sup>#</sup> | 6.3 ± 0.9 | 83.0 ± 10.1 | 6.9 ± 2.4 | n.d |
| Dehydrocostus lactone <sup>#</sup> | 1.4 ± 0.4 | 86.2 ± 4.1 | 5.8 ± 1.9 | 12.2 ± 0.5 |
| Eremanthin <sup>#</sup> | 1.9 ± 0.3 | 87.6 ± 9.9 | 7.3 ± 1.4 | 11.5 ± 0.2 |
| Bzn (20 μM) | 13.8 ± 2.9 | 76.9 ± 15.2 | >100 | n.d |

\* *Rumex crispus* root, <sup>†</sup> *Sium sisarum* root, <sup>‡</sup> *Rheum raphaniticum* root, <sup>#</sup> *Laurus nobilis* leaf

#### Impact of 9,10-anthracenedione substitutions on antichagasic effects in vitro

Emodin (**2**), which was representative for the anthraquinone cluster, moderately inhibited *T. cruzi* epimastigote growth and significantly inhibited parasite release in the cellular infection assay without being cytotoxic to host cells up to 100 µM (selectivity index of 200 (Table 3)). We therefore performed a preliminary structure-activity relationship (SAR) study on the 9,10-anthracendione (**1**) scaffold (Table 4). Both the inhibition of epimastigote proliferation and parasite release were measured. Our data indicate that the position of the hydroxyl groups was important for the antichagasic activity *in vitro*, with the trihydroxy-substituted derivatives being the most active (Table 5). The canonical anthraquinone (**1**) was ineffective against epimastigotes, and only very marginally active against cellular parasite release at 0.5 µM. Hydroxylation at R1/R2 (**4**) abolished this activity. Hydroxylation at R2/R6 (**9**) did not significantly improve activity of **1** in the infection assay. Likewise, the additional hydroxymethyl moiety at R3 found in aloe emodin (**11**) did not improve the activity in the infection assay but increased general cytotoxicity. Interestingly, hydroxylation at R1/R8 showed a trend towards general increase in toxicity towards epimastigotes (**2**, **8**, **10**-**12**). An acidic functional group (COOH) at R2 abolished the antichagasic activity as indicated by the lack of activity of rhein **(15)** and its clinically used anti-rheumatic prodrug diacerein (**16**) (see Table 5). The replacement of hydroxyl groups with methoxy and methyl groups generally led to a decrease in activity as shown by compounds **12** and **18**. Thus, 9,10-anthracenedione with hydroxyl groups at positions R1 and R3 or R4, as exemplified by compounds **2**, **3** and **6**, proved to be most active in the infection assay, but did not inhibit proliferation of epimastigotes. We also determined the long-term cytotoxicity (72 h incubation) towards host cells for the anthraquinones most effectively inhibiting *T. cruzi* parasite release in CHO cells (**2**,**3**,**6**,**14**). Noteworthy, all anthraquinones were cytotoxic only at higher micromolar concentrations, with the exception of quinizarin, which showed an IC_50_ value of 38 ± 16 µM. In this study, emodin (**2**) was the most potent antichagasic 9,10-anthracenedione in all stages of *T. cruzi,* followed by purpurin (**3**), which inhibited 50% (52.5 ± 14.4) of cellular parasite release at 0.5 µM.

**Table 4.** Chemical structures of natural and synthetic anthraquinones tested on *T. cruzi*.

| ID |  | R1 | R2 | R3 | R4 | R5 | R6 | R7 | R8 |
| --- | --- | --- | --- | --- | --- | --- | --- | --- | --- |
| 1 | Anthraquinone | H | H | H | H | H | H | H | H |
| 2 | Emodin | OH | H | OH | H | H | CH<sub>3</sub> | H | OH |
| 3 | Purpurin | OH | OH | H | OH | H | H | H | H |
| 4 | Alizarin | OH | OH | H | H | H | H | H | H |
| 5 | Alizarin RedS | OH | OH | SO<sub>2</sub>ONa | H | H | H | H | H |
| 6 | Quinizarin | OH | H | H | OH | H | H | H | H |
| 7 | Anthrarufin | OH | H | H | H | OH | H | H | H |
| 8 | Dantron | OH | H | H | H | H | H | H | OH |
| 9 | Anthraflavic acid | H | OH | H | H | H | OH | H | H |
| 10 | Chrysophanol | OH | H | CH<sub>3</sub> | H | H | H | H | OH |
| 11 | Aloe-emodin | OH | H | CH<sub>2</sub>OH | H | H | H | H | OH |
| 12 | Physcion | OH | H | CH<sub>3</sub>O | H | H | CH<sub>3</sub> | H | OH |
| 13 | 2-Hydroxy-3-methylantraquinone | H | OH | CH<sub>3</sub> | H | H | H | H | H |
| 14 | 2-Hydroxy-1-methylantraquinone | CH<sub>3</sub> | OH | H | H | H | H | H | H |
| 15 | Rhein | H | COOH | H | OH | OH | H | H | H |
| 16 | Diacerein | H | COOH | H | CH<sub>3</sub>COO | CH<sub>3</sub>COO | H | H | H |
| 17 | Disperse Red11 | NH<sub>2</sub> | CH<sub>3</sub>O | H | NH<sub>2</sub> | H | H | H | H |
| 18 | Aurantio obtusin | OH | CH<sub>3</sub>O | OH | H | H | CH<sub>3</sub> | OH | CH<sub>3</sub>O |

**Table 5.** *In vitro* antiproliferative activity of anthraquinones on *T. cruzi* epimastigotes (72 h) and trypomastigotes release (6 dpi). Compounds that inhibited at least 20% of parasite release in the trypomastigote infection assay at 0.5 µM were tested for antiproliferative effects on CHO host cells. Data shown are mean values ± SD of at least three independent experiments each performed in triplicates.

| ID | IC<sub>50</sub> epimastigotes<br>[μM] | % inhibition of parasite<br>release<br>at 0.5 μM | IC<sub>50</sub> CHO cells<br>[μM] |
| --- | --- | --- | --- |
| 1 | >50 | 17.8 ± 11.8 | n.d. |
| 2 | **14.1 ± 8.1** | **53.2 ± 24.3** | **>100** |
| 3 | >50 | 52.5 ± 14.4 | 59.5 ± 10.1 |
| 4 | >50 | 0 | n.d. |
| 5 | >50 | 0 | n.d. |
| 6 | >50 | 33.4 ± 20.1 | 38.1 ± 16.0 |
| 7 | >50 | 0 | n.d. |
| 8 | 5.3 ± 2.3 | 0 | n.d. |
| 9 | >50 | 22.4 ± 14.7 | 77.8 ± 1.7 |
| 10 | 6.6 ± 3.5 | 0 | n.d. |
| 11 | 14.5 ± 5.6 | 26.6 ± 21.9 | 42.0 ± 15.5 |
| 12 | 6.4 ± 3.7 | 18.1 ± 16.7 | n.d. |
| 13 | >50 | 0 | n.d. |
| 14 | >50 | 30.5 ± 7.9 | 88.9 ± 8.9 |
| 15 | >50 | 0 | n.d. |
| 16 | >50 | 0 | n.d. |
| 17 | 32.9 ± 13.9 | 0 | n.d. |
| 18 | >50 | 0 | n.d. |
| Benznidazole | 13.8 ± 2.9 | 0 | >100 |

## Discussion

### Biological profiling of plant extracts and assessment of the ethno-directed approach

In the present study we document the current knowledge about botanical drugs used to manage CD by the indigenous peoples Ayoreo, Chiquitano, Izoceño-Guaraní and Quechua in Bolivia. A major aim was to use this information for bioprospecting for anti-CD drugs. Previous studies reported better antimicrobial properties of extracts derived from botanical drugs selected based on popular uses related to microbial infections (59–61). Such studies lend credit to the ethno-directed approach in bioprospecting for specific bioactive metabolites. In order to challenge this approach, we tested a total of 775 EtOAc extracts from two independent botanical drug libraries generated from 115 taxa selected for their reported use against symptoms of CD in Bolivia and from 660 taxa described in *DMM*. Although the ethnomedical extraction generally proceeded with hot water infusions, we justify the use of EtOAc for extract preparation by the reduced extraction of polar and high molecular weight compounds, such as sugars and tannins, which would potentially interfere with the screening. Our findings indicate that the CD botanical drug library contains a significantly higher percentage of cytotoxic plant taxa. However, hit rates for selectively antichagasic plant extracts in the two libraries were not significantly different. The overall higher hit rate among the CD library was possibly due to non-specific cytotoxic effects, which could be conditioned by the ecological factors prevailing in the Chaco and Inter-Andean valleys. The extreme atmospheric and ecological conditions (altitude, extreme dryness and high temperature ranges) in these regions may favor the production of metabolites with broad-spectrum toxic or general antifeedant properties.

Among the indigenous peoples participating in this study, the biomedical concept of CD was only recently introduced and did not match with any existing traditional disease concept. This seems related to the fact that infection with *T. cruzi* shows a diffuse and varied disease pattern, and is asymptomatic in the majority of cases. We found that the majority of the botanical drug preparations intended for oral use (aqueous decoctions and infusions) were used to relieve symptoms associated with the chronic phase of CD, such as cardiac complications, fever and fatigue, and not for combatting the (invisible) parasites. In fact, most of the botanical drugs applied for CD related symptoms were also used for other therapeutic purposes involving inflammatory conditions (Table 2). We thus conclude that tangible ethnomedical concepts about CD were absent until recently and developed only during the last decades, which is in agreement with the cultural perception of *T. cruzi* vectors (41). This clearly hampers the application and selection of botanical drugs targeting *T. cruzi* parasitemia and its symptoms, and probably explains in part why the majority (> 80%) of the extracts derived from botanical drugs with reported use against CD and its symptoms were not active. The use of plant- and animal-based traditional medicine among the Chiquitano, Izoceño-Guaraní and Quechua was widespread and in agreement with previous reports that these people widely use traditional medicine despite the presence of Western healthcare (18, 62). Noteworthy, the Ayoreo did not treat CD at all, which is in agreement with their overall very limited use of herbal medicine. Chiquitano, Izoceño-Guaraní and Quechua research participants stated that they tried to manage CD with plant-based remedies because Western health care was limited, and chemotherapy not accessible during the chronic stage of CD. Another study from a different region in Bolivia reported a similar situation (34). Asteraceae was the most dominant family of plants usind for CD, likely due to their overall abundance and species richness. In general, Asteraceae are over-proportionally represented in medical florae (63–65), and this may be linked to the high diversity of bioactive secondary metabolites in the family (63). Quechua informants showed a low consensus regarding the species to be used in the treatment of CD with only one species mentioned three times. A higher consensus was found among the Chiquitano and Izoceño-Guaraní participants (estimated Trotter and Logan Informants’ consensus: Fic > 0.8) with relatively few species being used by a large proportion of participants. Among the Izoceño-Guaraní *S. chloroclada* (referred to as lanza lanza, mbuijare or retama) was clearly the most important species for treating CD, with a share of 48% of total use-reports. Interestingly, *S. chloroclada* belonged to a phylogenetic cluster that showed no or little inhibition of epimastigote proliferation in the pre-screening (Fig. 4), but exhibited significant antichagasic effects in the parasite release assay (79.5% inhibition at 15 μg/mL by EtOAc extract of flower). This discrepancy was also observed with some anthraquinones (purpurin and quinizarin). Since the extract likely contains glycosides that may be hydrolyzed by CHO cells but not by epimastigotes, we cannot exclude metabolic changes induced by host cells. The genus *Senna* (syn. *Cassia*) is known to contain anthraquinone, dianthrone and naphthol glycosides (66). A preliminary ESI-MS scan analysis and the Borntrager reaction confirmed the presence of anthraquinones (physicon and chrysophanol) in this botanical drug (supplementary Fig. S1). To date, no in-depth phytochemical study is available on *S. chloroclada*, and follow-up studies are planned in our laboratory. *T. aurea* bark as its Spanish vernacular name “paratodo” indicates, was used for numerous diseases, in agreement with a previous study (67). The introduced species *A. zerumbet* and *C. citratus* are well known in South America for the treatment of cardiovascular diseases (67–70). An apparently specific medicinal indication for CD was also reported for several species mentioned by our research participants, such as *Aloysia citridora*, *Baccharis genistelloides*, *Bixa orellana*, *Dysphania ambrosioides*, *Handroanthus impetiginosus*, *L. usitatissimum*, *Plantago major*, *Ruta chalepensis*, *S. chloroclada, S. junceum* and *Schinus molle* (17,34,35,71–76). None of the EtOAc extracts obtained from botanical drugs of these species were significantly and selectively toxic against *T. cruzi* in our infection assay with the exception of *S. chloroclada* (flowers) inhibiting parasite release > 50% *in vitro* at 15 μg/mL (Suppl. Table S1) but not showing any effect on epimastigotes.

The extract obtained from the aerial parts of *A. buniifolius* (Fig. 3) had a SI > 20 and was the other noteworthy hit obtained with the CD informed library. It fully inhibited epimastigote proliferation at 25 μg/mL and parasite release at 15 μg/mL. Its antichagasic and antileishmanial activity was previously reported for plant material collected in Argentina, and the flavonoid santin was thought to be the active compound (77). However, santin was only moderately active against *T. cruzi* epimastigotes and trypomastigotes (IC_50_ values > 30 µM). Clearly, the presence of additional antichagasic metabolites should not be excluded, and an in-depth phytochemical investigation of *A. buniifolius* is warranted. The extract of seeds of a *Pterodon* sp. showed significant and apparently specific antitrypanosomal effects. Whether the very common diterpene alcohol geranylgeraniol previously identified as the major antichagasic component in *Pterodon pubescens* seeds (78) is responsible for this specific antitrypanosomal activity remains to be clarified.

The screening of the *DMM* extract library resulted in 23 extracts with selective parasite toxicity in the *T. cruzi* release assay. Of these extracts, eight were from Apiaceae and seven from Asteraceae showing taxonomic parallels with the CD-informed library. In both libraries, Asteraceae was the largest family. Asteraceae are known to frequently contain SLs with antiprotozoal properties (79, 80) (*vide infra*). It is possible that the use of SL containing botanical drugs could therefore represent an ethnopharmacological strategy to reduce parasitemia in CD. The 660 extracts representing the *DMM* library were obtained from 389 different plant species. The largest part are from the Mediterranean basin, but central European species and exotic herbal drugs imported from Africa, Arabia, Central Asia, Himalaya and the Indo-Malayan region are also included. The families with the highest share of botanical drugs and species are Apiaceae with 69 botanical drugs from 37 spp., Asteraceae with 51 botanical drugs from 33 spp., Rosaceae with 37 botanical drugs from 18 spp., Lamiaceae with 33 botanical drugs from 26 spp. and Fabaceae with 29 botanical drugs from 22 spp. This pattern reflects the overall taxonomic composition of *DMM* for which a total of 536 plant taxa representing 924 botanical drugs were identified and recommended for 5314 medical applications (40). The medical categories with the most therapeutic uses are dermatology (1216), gastroenterology (805), gynecology (615), urology (437), respiratory system (374) and neurology (269). The most frequently mentioned parasite treatments are related to lice, scabies and tapeworms but also applications for malaria causing infections with *Plasmodium* were described (tertian and quartan fever) such as all parts of *Anchusa* sp., seed and leaves of *Bituminaria bituminosa*, root of *Dipsacus fullonum*, seeds of *Heliotropium europaeum*, root of *Plantago* sp., herb of *Potentilla reptans* and the herb of *Verbena officinalis* (40). With respect to the overall Mediterranean flora the Apiaceae and Rosaceae appear to be overrepresented in this library, while the frequency of Asteraceae and Lamiaceae appears to be rather consistent with the overall species diversity and abundance. Apiaceae fruits were frequently used as antidotes, and their resins for neurological and musculoskeletal problems. Considering the size of plant families, the Fabaceae seem underrepresented, while Poaceae, Caryophyllaceae and Orchidaceae are clearly underrepresented in the *DMM* library (Suppl. Table 2S; (40, 50)). With respect to the treatment of fevers and parasites, the active extracts from plants belonging to the ‘cumarin cluster’, such as the herb of rue (*R. chalepensis*) were recommended in *DMM* for the internal use of the treatment of tremor and shivering prior to fever attacks while the seeds of *Seseli tortuosum* against fevers in general. The hits belonging to the ‘anthraquinone cluster’, such as the roots of *Rumex* species and rhubarb (*Rheum* spp.) were recommended for the treatments of scabies and fevers, respectively. For *L. officinale*, *L. nobilis* and *S. sisarum* no uses related to fever and parasites are recorded in *DMM* (40).

### Microfractionation and isolation of bioactive principles of selected antichagasic plant taxa from the DMM library

For the isolation of active compounds from the *DMM* library we selected extracts pertaining to three different phylogenetic groups, with taxa whose plant material was readily accessible. A major phylogenetic hotspot showing apparent antitrypanosomal selectivity was the anthraquinone cluster with the rhizomes of *R. crispus* (curly dock) and *R. rhaponticum* (rhapontic rhubarb) from the Polygonaceae family. The anthraquinones emodin and chrysophanol were identified in *R. crispus* and served as a basis for the subsequent preliminary SAR study (*vide infra*). Since the naphthoquinone derivative nepodin with known antimalarial activity (75) was isolated together with anthraquinones from *R. crispus* and likewise inhibited epimastigote proliferation, the potency of the extract may reflect additive effects. In general, naphthoquinones have been studied extensively in *T. cruzi* epimastigotes (81, 82) without leading to a successful translation to clinical trials (83). Extracts with antitrypanosomal activity from plants with distinct phylogenetic positions were those from *L. nobilis* (bay) (Lauraceae) and *S. sisarum* (skirret; Apiaceae). *L. nobilis* leaves contain a range of SLs which may explain the significant antichagasic effects of its extract in epimastigotes. Numerous studies have addressed the selective versus non-selective antitrypanosomal effects of SLs which are also widely present in Asteraceae (84–86). SLs like cynaropicrin and others can act as electrophiles and form adducts with biological nucleophiles, such as trypanothione, the parasitic equivalent of glutathione in mammalian cells (26, 87). The reason why *L. nobilis* (leaves) was ineffective in the infection assay could be due to the SLs reacting with thiols in host cells, e.g. glutathione, without reaching the parasite. It has been shown that CHO cells can produce glutathione upon stress (88). Thus, SLs undergoing a Michael-type addition with thiols are likely poorly bioavailable to infected tissues as they are detoxified by glutathione. Although the polyacetylene falcarindiol from *S. sisarum* had no inhibitory effect on *T. cruzi*, we cannot exclude the possibility that other polyacetylenes present in *S. sisarum* root may be more potent as indicated by the activity profile of the extract.

### SAR study of anthraquinones as antichagasic natural products in vitro

Anthraquinones are condensed aromatic hydrocarbons found in different plant species known for their medical and dye applications (89). In Western pharmacopoeias, anthraquinone containing botanical drugs, such as *Rhamnus* spp. (Frangulae cotex and Rhamni purshiani cortex) or *Rheum officinale* (Rhei radix) are used as laxatives. The 9,10-anthracenedione scaffold, if bioavailable to the infected cells, could interfere with the parasite redox system as this scaffold can mediate the production of hydrogen peroxide or reactive oxygen species (ROS) via oxygen reduction *in situ* (90). Anthraquinones have been shown to interfere with redox reactions in cells (91). ROS generation is also the postulated mode of action of the approved antichagasic drug benznidazole (92). However, further studies will be necessary to uncover the underlying mode of action. To elaborate on our natural product drug discovery approach, the most active antichagasic secondary metabolite emodin (1,3,8-trihydroxy-6-methylanthracene-9, 10-dione) led us to explore in more detail the SAR of differentially substituted 9,10-anthracenediones. Trihydroxylated anthraquinones have already been shown to be trypanocidal (*vide infra*), but this study provides a first preliminary SAR study on 9,10-anthracenediones for both *T. cruzi* epimastigote and parasite release from trypomastigote infected CHO cells. Different anthraquinones have been shown to exert moderate to good antimalarial, antibacterial and antiviral effects *in vitro* at low micromolar concentrations (93). In our study, emodin showed specific submicromolar inhibition of parasite release in CHO cells. Emodin was previously only tested on epimastigotes at high micromolar concentrations (94) where it inhibited casein kinase 1 with an IC_50_ value of 130 µM (95), thus significantly less potent than in the trypomastigote infection assay shown here. Other reported effects of this natural product include antiinflammatory, antiosteoporosis, and antidepressant effects (93). Purpurin, a natural trihydroxylated anthraquinone, inhibited parasite release and showed a selectivity index > 100. Rather unexpectedly, it did not inhibit epimastigote proliferation, which may be due to differences in pH between lysosomes and culture medium related to the trypanosomal uptake of anthraquinones. Purpurin has previously been shown to inhibit blood-stream trypomastigotes (96), but was not studied in cellular infection assays. Since anthraquinone glycosides in plant extracts can be deglycosylated and reduced to anthrones and anthranoles by gut bacteria (97), resulting in potent laxative effects, the systemic application of these compounds can be challenging. However, as exemplified by the drug diacerin (prodrug of rhein), low micromolar plasma concentrations can be achieved (98). Unfortunately, rhein showed no antitrypanosomal effects in our assays, possibly due to the carboxylic acid at C2. It is noteworthy that the most important botanical drug used in the context of CD among the Guaraní also contains anthraquinones, thus giving strong support to the anthraquinone cluster being of potential relevance in the treatment of CD.

Overall, the comparative phylobioactivity-guided screening for in vitro antichagasic activity is an enabling tool in the discovery of natural products that may have a potential in CD drug development. The comparative profiling was further suitable for a pharmacological validation of a library with botanical drugs currently used in the context of CD in Bolivia, thus challenging the ethno-directed CD bioprospecting approach. Moreover, our study led to the identification of biologically significant antichagasic phylogenetic hotspots that will serve for subsequent phytochemical investigations.

## Acknowledgements

This study was funded by the EU Marie Curie Actions Initial Training Network (ITN) MedPlant (medplant.eu). We would like to acknowledge the Guaraní, Ayoreo, Chiquitano and Quechua informants who agreed to participate in the research, and the members of the communities for their hospitality and interest. Rossy Chávez de Michel and Stephan Beck from the National Herbarium Bolivia (LPB) identified the herbarium specimens. We thank Yonny Flores, Alberto Giménez and Efrain Salamanca (Universidad Mayor de San Andres) for technical assistance in the lab in Bolivia and for introducing the handling of *T. cruzi*. We acknowledge Matthias Rubin (University of Bern) for technical assistance in the lab and Peter Staub (Unica) for collecting *DMM* plant material and Laura Casu (Unica) for production of extracts. We thank Peter Bütikofer (University of Bern) for sharing the *T. brucei* strain and giving advice on its culturing.

